# yEvo: a modular eukaryotic genetics and evolution research experience for high school students

**DOI:** 10.1101/2022.05.26.493490

**Authors:** M. Bryce Taylor, Alexa R. Warwick, Ryan Skophammer, Josephine M. Boyer, Renee C. Geck, Kristin Gunkelman, Margaux Walson, Paul A. Rowley, Maitreya J. Dunham

**Affiliations:** Department of Genome Sciences, University of Washington, Seattle, WA 98195; Program in Biology, Loras College, Dubuque, IA 52001; Department of Fisheries and Wildlife, Michigan State University, East Lansing, MI 48824; Westridge School, Pasadena, CA 91105; Department of Biological Sciences, University of Idaho, Moscow, ID 83844

## Abstract

Microbial experimental evolution paired with whole-genome sequencing allows researchers to observe evolutionary processes in real-time. The resources for carrying out and analyzing microbial evolution experiments have become more accessible. It is now possible to expand these studies beyond the research laboratory and into the classroom. We have developed a series of five connected and standards-aligned yeast evolution laboratory modules, called “yEvo,” for high school biology students. The modules have been designed to enable students to take agency in answering open-ended research questions. In Module 1, students evolve the baker’s yeast *Saccharomyces cerevisiae* to tolerate an over-the-counter antifungal drug, and in subsequent modules, investigate how evolved yeasts adapted to this stressful condition at both the phenotype and genotype levels. Pre- and post-surveys from 72 students at two different schools and one-on-one interviews with students and teachers were used to assess our program goals to iteratively improve these modules over three years. We also measured changes in student conceptions of mutation and evolution, confidence in scientific practices, and interest in STEM and biology careers. Students who participated in our experimental evolution module showed improvements in activity-specific concepts, including the importance of variation in evolution and the random nature of mutation. They additionally reported increased confidence in their ability to design a valid biology experiment. Student experimental data replicated literature findings on mechanisms of clotrimazole resistance and has led to new insights into this phenomenon. This collaborative endeavor will serve as a model for other university researchers and K-16 classrooms interested in engaging in open-ended research questions using yeast as a model system.

## Introduction

A growing movement in STEM education aims to incorporate research experiences into the K-16 classroom (Sadler et al. 2010). These efforts have taken on many names, including course-based research experiences (CREs), student-scientist or school-scientist partnerships (SSPs; (Clendening 2004)), and course-based undergraduate research experiences (CUREs; (Auchincloss et al. 2014; Krim et al. 2019)). Research experiences have improved students’ confidence, grasp of concepts, and interest in STEM careers (Auchincloss et al. 2014; Krim et al. 2019; Indorf 2019; Hunt 2021). They may help with recruiting and retaining women and students from underrepresented backgrounds (Bangera 2014; Hunt 2021). Additionally, participation in research can have positive impacts on teachers’ professional development (Silverstein 2009).

The value of research experiences has led to a growing body of research and implementations at colleges and universities (see CUREnet https://serc.carleton.edu/curenet/index.html). Compared to the college level, published examples of research experiences have been more limited in high school (Tanner et al. 2003). This paucity is in spite of the fact that at the K-12 level, these efforts are in alignment with the Next Generation Science Standards (NGSS) (NGSS 2013), which have made participation in the scientific process through inquiry-based activities a central organizing principle in curriculum design.

Microbial experimental evolution, sometimes referred to as adaptive laboratory evolution, is appealing for classroom-based research activities because these experiments have relatively low resource requirements and can be carried out in a few days or weeks. Experimental evolution has been successfully utilized in teaching modules for K-12 and college courses (Cooper 2019; Ratcliff 2014; Smith 2016; Bennett 2021). However, we have identified very few resources for conducting evolution research activities outside of college and university settings (example education conference report: (Laursen et a. 2007)). It is worthwhile to develop resources to facilitate these activities since experimental evolution can connect concepts in evolution, cell biology, genetics, medicine, and biotechnology. These connections are also relevant to the NGSS emphasis on “cross-cutting concepts” that demonstrate commonalities between disciplines that are often taught separately.

In the experimental evolution framework, populations of organisms, most commonly microbes, are propagated under suboptimal conditions that restrict growth, such as elevated temperatures, exposure to drugs, or nutrient limitation. Over time, this will select for mutant organisms that are better adapted to such environments (reviewed in Kawecki 2012; Lang 2014; McDonald 2019). As a population of microbes is propagated in a suboptimal growth environment, rare mutations that lead to enhanced growth rise in frequency due to natural selection until they constitute an increased fraction of the microbial population. This change results in phenotypic differences at the population level (i.e., increased growth), allowing the observation of the process of evolution in real-time. Whole-genome sequencing of mutant organisms isolated from these experiments can provide insight into the genetic and molecular changes that lead to adaptation to a specific selective pressure (Long 2015; Payen 2016; Bruger 2018). Traditionally this experimental paradigm has addressed basic evolutionary questions, such as the speed, dynamics, and limits of adaptation (e.g., Kawecki 2012; Lenski 2017; McDonald 2019). It is increasingly implemented in applied contexts as a form of domestication to isolate microbes adapted to industrial settings (Giannakou et al. 2020; Sandberg 2019; Lee 2020).

For several reasons, the budding yeast *Saccharomyces cerevisiae* is a popular species for experimental evolution and an attractive microbial model system for high school students (Duina 2014). (1) *S. cerevisiae* can be grown safely and easily within a classroom environment without specialized equipment. (2) It is one of the most well-characterized organisms from a molecular and genetic standpoint due to extensive work both in laboratories and in broadly-familiar settings such as baking and fermentation. (3) Its small, highly-annotated genome, ease of phenotyping, and short generation time make it ideal for experimental evolution studies. (4) It is a key model organism that has yielded many discoveries about fundamental biological principles. It is also an important industrial organism used to produce proteins and small molecules for pharmaceutical and biotechnology applications. (5) Thousands of academic and industrial researchers are working in this active field in both basic and applied research (e.g., Botstein 2011; Lee 2020).

To address the need for authentic research experiences in high school that integrate learning objectives in evolution, cell biology, and genetics, we developed protocols for the experimental evolution of *S. cerevisiae* in a high school classroom setting, named “yEvo” (yeast Evolution). These experiments utilize strains of yeast engineered to express vibrant pigments (**Figure 1**). These strains provide many technical advantages (**Supplemental Text 1**), including a simple means to monitor for culture contamination or mislabeling, which are consistent issues in microbiology experiments. Our protocols were developed in close collaboration with high school teachers to ensure that they would be compatible with classroom learning objectives, teacher interests, and practical constraints. We leveraged the flexibility of our experimental system to develop multiple versions of our evolution protocol to suit a variety of time and resource availabilities. Our protocols differ from previous evolution teaching modules in that our experimental question is open-ended, and the classroom projects have the opportunity to contribute to an ongoing research program. Key features in our design include opportunities for agency (choosing aspects of experimental conditions for selection), competition (experiments to compare the fitness of evolved yeast), and collaboration (working together in teams, performing a literature search based on student mutation data).

**Figure 1.**
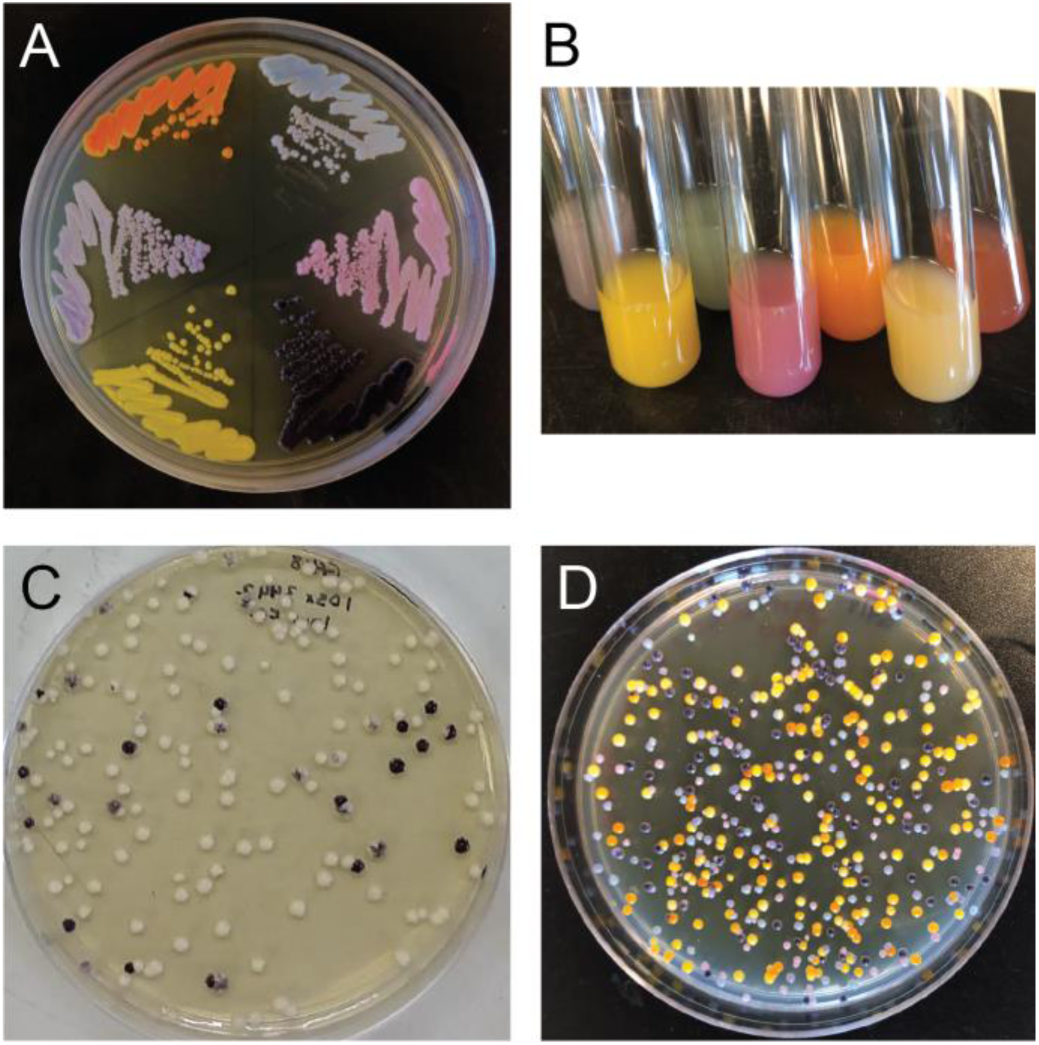
Yeast expressing different color pigments grown (A) on solid media and (B) in liquid culture. (C) Two or (D) six yeast strains expressing different pigments can be differentiated after co-culture.

Our initial yEvo protocols tasked students with identifying molecular factors that allow *S. cerevisiae* to become resistant to an over-the-counter azole-class antifungal drug (clotrimazole) (Allen 2015; Shafiei 2020). This topic is highly relevant because antifungal resistance among pathogenic fungi is a growing global health threat and also threatens food security (e.g. (Berkow 2017; Fisher 2018)). Research into the genetic basis of this trait has allowed medical researchers to develop new approaches to treat drug-resistant pathogens (e.g., Cowen 2009 PNAS)) and to predict which treatments are most likely to work for each patient’s infection (e.g. (Berkow 2017; Cowen 2015). *S. cerevisiae* is a safer alternative to experiments with pathogenic fungi as it cannot cause disease in healthy people yet shares many of the genes involved in antifungal resistance with pathogenic fungi.

This paper presents our yEvo teaching protocols, organized into five stand-alone modules (**Figure 2**) that can connect to various topics in standard biology curricula (e.g., College Board and NGSS). We discuss reflections on our design and iteration process over the first three years of implementation, which were guided by interviews and surveys with students and teachers. Our survey evaluations measured changes in disciplinary knowledge, student STEM career interest, confidence with scientific practices, and general engagement with the material in three high school teachers’ classrooms in two different schools. Critically, students reported increased confidence in their ability to design a valid biology experiment and expressed an increased interest in a STEM or biology career due to participating in yEvo. The student experimental data replicated literature findings on mechanisms of clotrimazole resistance and led to new insights into this phenomenon (Taylor 2021). Teachers felt that these investigations with college-level collaborators removed barriers to course-based research. This collaborative endeavor will serve as a model for other university researchers and high school teachers interested in engaging K-12 students in authentic research experiences.

**Figure 2.**
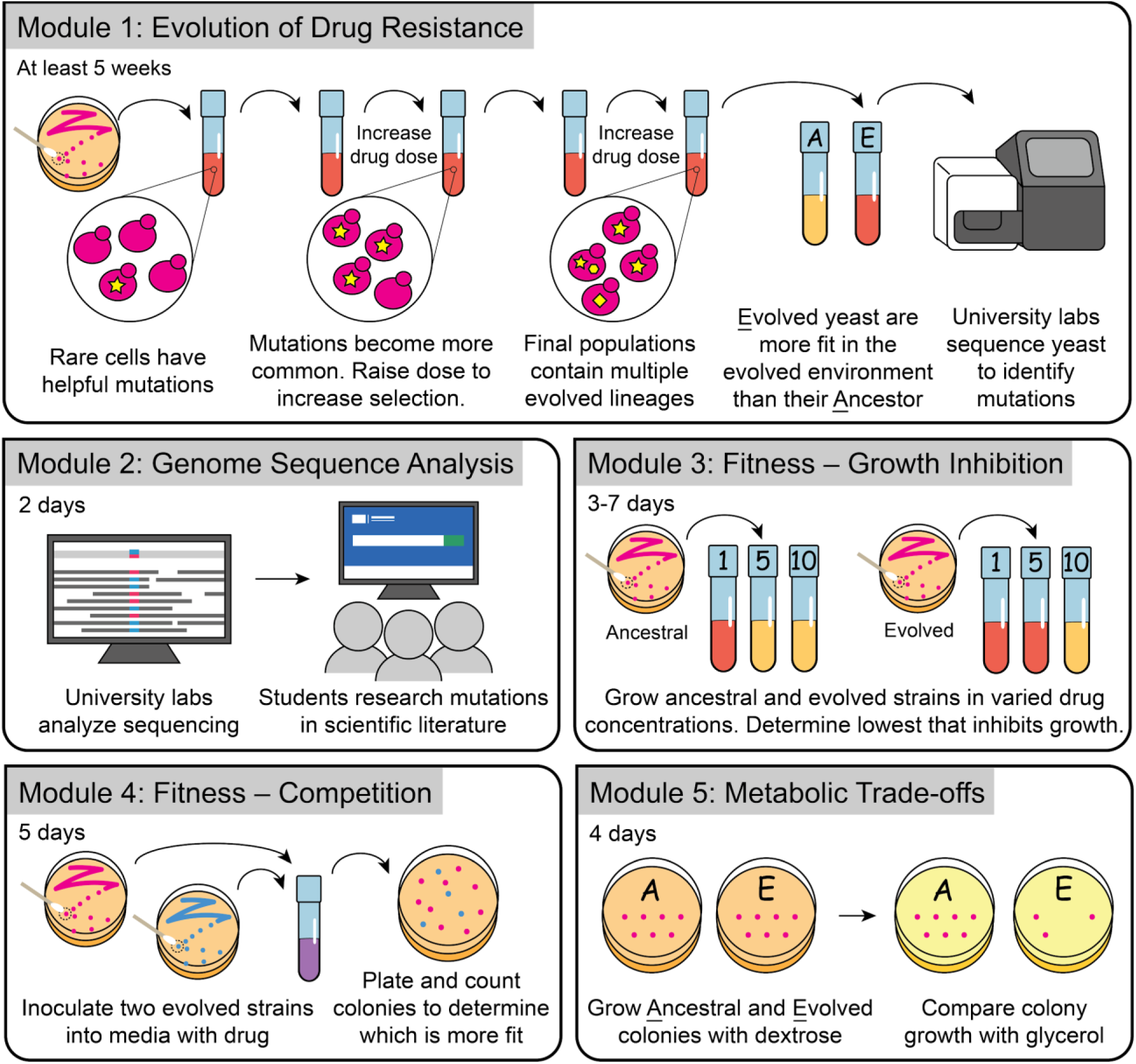
Overview of yEvo Modules 1-5.

## Methods

### Motivation

The goal of yEvo is to involve students in an authentic research experience that connects genetics, cellular and molecular biology, and organism-level phenotypes in an evolutionary context (**Figure 2**). As such, yEvo addresses complex and often abstract concepts, such as randomness and temporal scales, with which students frequently struggle (Dougherty 2011; Tibell 2017). We developed yEvo in collaboration with teachers at two US schools, a private school in California and a public school in Idaho, to meet the needs of the teachers and their students. The module activities were tailored for the time and resources available to each teacher, which resulted in some differences among the focal classrooms in this study (**Figure 3**). The modularity of yEvo allows teachers flexibility in including it in their curriculum. Modules are also designed to touch on many topics in a standard biology curriculum. During the five modules, students select for yeast mutants that are more resistant to clotrimazole, examine genome sequencing data to identify mutations that may be responsible for this resistance phenotype, and use cellular and molecular models to contextualize how their mutations may be connected to the resistance phenotype. For each module, lab skills and suggested discussion topics with standards can be found in **Table 1**. Additional information on our design philosophy can be found in **Supplemental Text 1**.

**Table 1.**
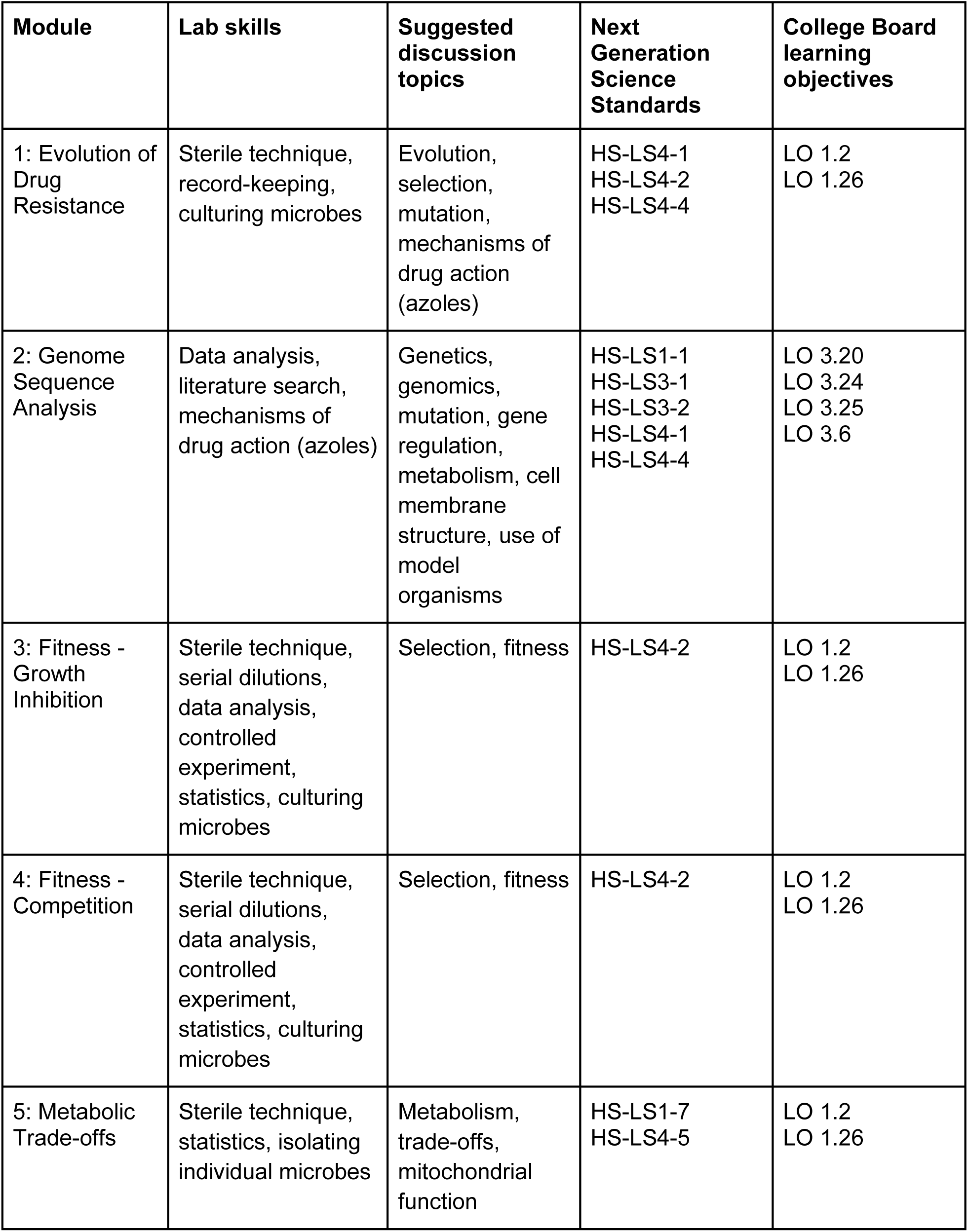
Module ties to standards.

**Figure 3.**
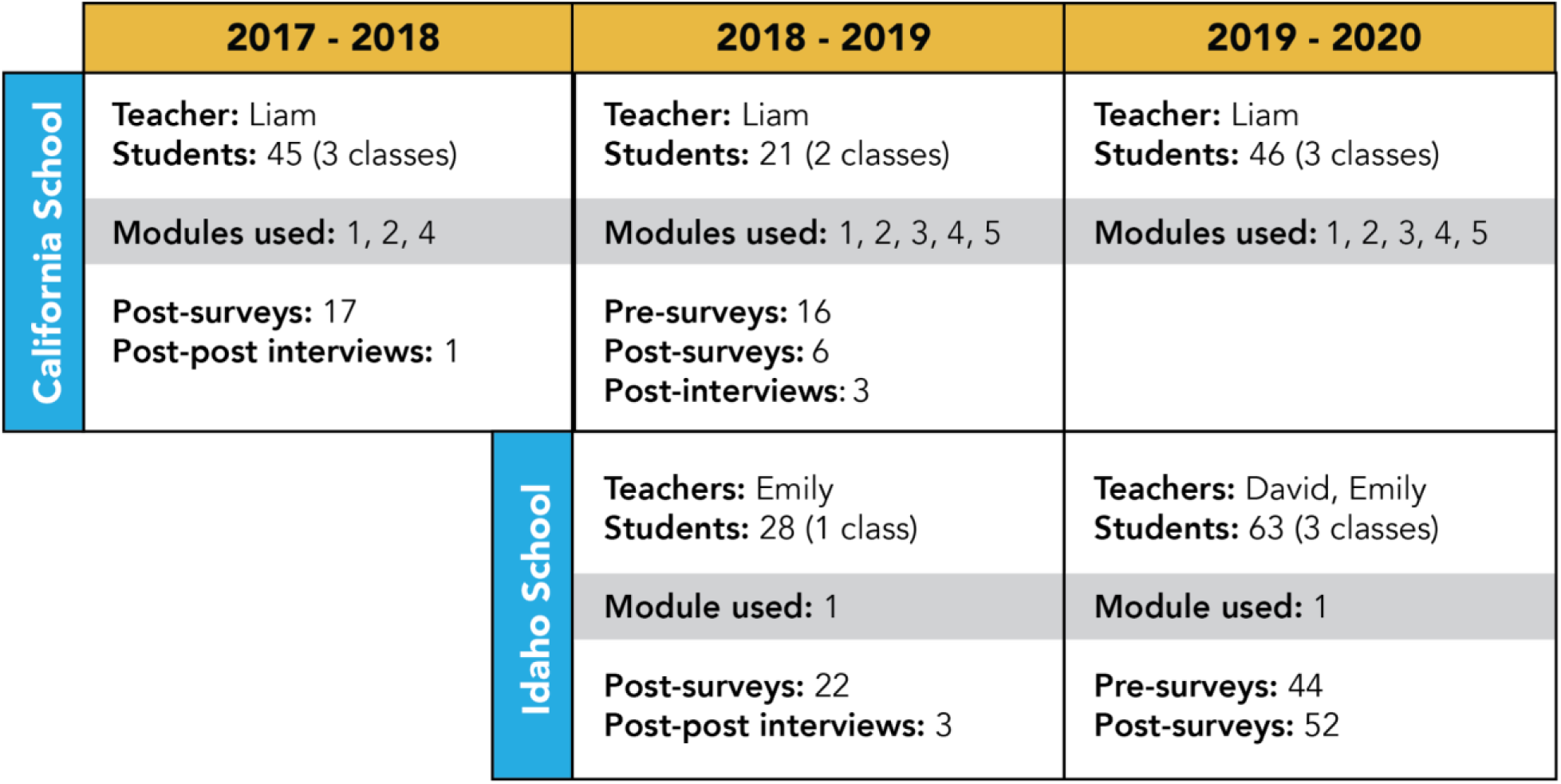
Timeline of module implementation for each school/teacher, number of students involved, and evaluation collected.

Briefly, in Module 1 (‘Evolution of Drug Resistance’) students grew yeast in the presence of an over-the-counter antifungal azole drug (FungiCure; active ingredient clotrimazole) for several weeks (**Figure 2**). Students transferred their yeast into new drug-containing medium at regular intervals and increased the drug dosage as they observed improved growth. The length and frequency of interaction with these experiments were flexible to classroom time constraints. Resistance phenotypes can be reliably observed after five transfers, which can be performed at intervals of 2 days or up to 2 weeks. Students carried out evolution experiments for 7 to 34 weeks, depending on the year and the classroom, using one of two protocols (**Supplemental Text 2**). We isolated clones with increased clotrimazole resistance from some experiments as early as two weeks (Taylor 2021).

The remaining modules allow students to investigate mechanisms of adaptation in their yeast from Module 1 or in yeast isolated from prior classrooms that completed this module. Module 2 explores the genetics of evolution. We have determined the genome sequence of 99 clones of yeast from student experiments and identified mutations that occurred during these experiments. Students use freely-accessible online databases such as the *Saccharomyces* Genome Database (yeastgenome.org) and NCBI BLAST (https://blast.ncbi.nlm.nih.gov/Blast.cgi) to learn about these mutant genes (**Figure 2**). Modules 3 and 4 center around measuring adaptive phenotypes in evolved yeast (**Figure 2**). Module 4 utilizes a competition experiment in which yeast expressing different pigments are mixed and grown in the presence of clotrimazole and then plated on an agar-based medium. If one strain in the mixture has a higher fitness, it will produce more viable cells, which can be observed by counting colored colonies on the agar surface. Finally, Module 5 investigates a metabolic trade-off (**Figure 2**). Evolved yeasts that are resistant to clotrimazole frequently lose the ability to carry out cellular respiration, which provides a growth advantage in the presence of clotrimazole. However, these resistant mutants cannot grow on a medium containing a non-fermentable carbon source, demonstrating that some evolved characteristics can be detrimental in the wrong environment (a trade-off). More details about each module and its protocols can be found in (**Supplemental Texts 1-6**) and on our website yEvo.org. For access to data, see our publication on mechanisms of azole resistance (Taylor 2021).

### Teacher and student characteristics

The three focal teachers (referred to here as Liam, Emily, and David) are all experienced teachers who have been in their current positions for over eight years. All teachers are white, one female and two male. Liam holds a Ph.D. in cellular and molecular biology, and Emily has a Master’s in wildlife biology. Liam teaches AP Biology at an all-female school in California, and Emily and David both teach 10th grade Honors Biology at a mixed-gender school in Idaho. As part of our pre/post surveys (described below), we asked students to respond about their race/ethnicity, age, and primary language spoken at home. Data reported here are from 14 of Liam’s students (all female) who responded to our surveys in 2018-2019, and 63 of David and Emily’s students (50-52% female) who responded to surveys in 2019-2020 (**Table 2**). Across both schools, most students spoke English at home; a few students under each teacher spoke other languages at home (**Table 2**). Liam’s students were between 15-17 years old, and 64.3% identified as non-white (**Table 2**). On average, the Idaho students were one year younger than Liam’s students, and between 16.2-28.6% identified as non-white (**Table 2**). All California students had taken a previous 9th grade biology course, and about two-thirds (69%) had previous lab course experience. A small number of students (3/16 on the 2018-2019 pre-lab survey) reported having worked in a professional lab or clinic setting. This was the first biology-focused science course for Idaho students, though they had some biology exposure from a 7th grade life science course.

**Table 2.** Characteristics of participants include the number of unique students with data collected from pre/post surveys for each of the three teachers, percent of female students, percent of students who identified with a non-white race/ethnicity (could also identify as white / multi-racial), the mean age of students (if they responded to both pre- and post-survey we used their age at post), and the primary language spoken at home. Student gender and racial demographics are representative of the school as a whole based on publicly-available demographic records.

| Teacher (State) | No. of Students | %Female | %Non-White | Age (mean) | Language(s) spoken at home |
| --- | --- | --- | --- | --- | --- |
| Emily (ID) | 21 | 52.4% | 28.6% | 15.33 | 19 English<br>1 Chinese<br>1 Nepali |
| David (ID) | 37 | 50.0% | 16.2% | 15.69 | 36 English<br>1 English/Mandarin |
| Liam (CA) | 14 | 100.0% | 64.3% | 16.35 | 11 English<br>1 English/Spanish<br>1 English/Spanish/German<br>1 Arabic |

### yEvo development and implementation

Modules 1, 2, and 4 were developed and piloted during the California school’s 2017-2018 school year (**Figure 3**). Modules 3 and 5 were developed during the 2018-2019 school year and piloted at the California school (**Figure 3**). All protocols are based on standard research lab practice but modified in collaboration with classroom teachers to fit their needs. The Module 1 protocol was modified to fit the class structure at the Idaho school by having students transfer yeasts to a new growth medium once per week and carry out the Module 1 experiment for 15 weeks. Detailed protocols can be found in **Supplemental Texts 1-6**. Teachers incorporated lab activities with consultation from researchers but followed their existing classroom plans. In addition to the modules described, some students from these classes participated in other research-related activities and field trips (e.g., additional experiments, conference presentations, visits to researcher labs) before or after the modules were implemented.

### Survey development

We developed an online survey using published questions (Jeffery 2016; Richards et al. 2017) and novel yEvo-specific questions related to the modules. Our goals were to evaluate (1) how students conceptualized topics introduced through yEvo, (2) which aspects of yEvo students liked and disliked, (3) how yEvo impacted students’ confidence in their ability to perform scientific investigations, and (4) changes in students’ interest in STEM and biology-related careers. Interview questions were developed to follow up with student responses to survey questions via remote interviews (on Zoom). Prior to any module activities, parents/guardians were sent a handout with information about the study and a form to return if they did not consent to share their child’s data for the study (passive consent). Students also assented to share their data through either a hardcopy or online form. All research methods were submitted and determined to qualify for exempt status by the University of Washington (IRB #00003148).

### Survey data collection

The modules and surveys were implemented across high school classrooms from 2017 through 2020 (**Figure 3**). In year one, we only administered a post-lab survey in California to test the survey questions on this new population and receive initial student feedback to make adjustments to modules for year two. In year two we administered both pre- and post-lab surveys at both schools and conducted semi-structured interviews with a subset of California students from both school years. Because of differences in module usage resulting from school closure during the COVID-19 pandemic, we report our results for the two teachers in Idaho separately unless noted.

We also collected the AP Biology test scores from 2014-2019 classes at the California school. We calculated a weighted score for the class using the formula 1/2(mean multiple-choice score)+1/2(mean free response question score). We additionally calculated a global average for all who took this exam. The values shown are the differences between those scores.

### Analysis of student evaluations

Quantitative survey responses were analyzed by comparing the teacher group averages and individual student changes in the pre- and post-survey using t-tests. To directly compare pre- and post-changes, we used only paired pre-post responses for some analyses. We used averages of all collected responses for other analyses, even if a student did not complete both surveys. Short-answer survey questions were coded for response themes using qualitative content analysis, primarily a summative approach (Hsieh and Shannon 2005). We developed a coding scheme to pull key terms or phrases from open-ended responses to questions 1-4 and 6 (**Table 3**) by reviewing learning objectives based on NGSS, drafting an ideal correct response, and then reading a subset of student responses for emergent themes. Thus, we also included three codes intended to capture responses indicative of an incomplete understanding or misconception (‘make’ in Q1; ‘vague adapt’ and ‘naive’ in Q3). We include a summary of code names and descriptions (**Supplemental Tables 1-4 and 6**). Student responses were coded in a binary system, with ‘0’ meaning the student did not include the key term or concept in their response and ‘1’ meaning the student did include the key term or concept in their explanation. Student responses could include zero or multiple codes for a single question. In contrast to the other questions, student responses to question 5 (**Table 3**) were evaluated on a 0-3 point rubric for correctness, summarized in (**Supplemental Table 5**). All codes and the scoring rubric were iteratively developed among three co-authors (multiple rounds of inter-rater coding comparisons) and coded blind with regard to student classroom and pre or post. For analysis, most Likert-response questions were converted to a numerical scale where Strongly Disagree = 1, Disagree = 2, Neutral = 3, Agree = 4, and Strongly Agree = 5.

**Table 3.** Survey questions, analysis method used, and location of additional information in supplemental tables.

| Question | Analysis | Code description | Results |
| --- | --- | --- | --- |
| 1. What is a gene? | Coded for key terms / concepts and alternative conceptions | Supplemental Table 1 | Table 4; Figure 4 |
| 2. What is a mutation? | Coded for key terms / concepts and alternative conceptions | Supplemental Table 2 | Table 5 |
| 3. How would you describe evolution? | Coded for key terms / concepts and alternative conceptions | Supplemental Table 3 | Table 6; Figure 4 |
| 4. What role do mutations play in evolution? | Coded for key terms / concepts and alternative conceptions | Supplemental Table 4 | Table 7; Figure 4 |
| 5. How would you explain antibiotic resistance to a fellow student in this class? | Scored based on correctness (0, 1, 2) | Supplemental Table 5 | Figure 5; Supplemental Figure 2 |
| 6. Individual microbes develop mutations in order to become resistant to an antibiotic and survive. | Compared level of agreement with open-ended explanation. Coded explanation for key terms / concepts and alternative conceptions. | Supplemental Table 6 | Table 8; Figure 6; Supplemental Figure 3 |

## Results

### Development of yEvo modules in California classes 2017-2018

The first implementations of yEvo were at the California school during the 2017-2018 academic year, in which students participated in Modules 1, 2, and 4, which included performing the evolution experiments, analyzing the mutations in the evolved strains, and competing evolved strains to measure fitness (**Figure 3**). A total of 17 students completed a post-lab survey. Overall, most students (16/17) stated they were willing to participate in yEvo again because it was “fun.” One student wrote, “After lots of years of learning science exclusively in a classroom, it was fun to feel like we were doing ‘real’ science and seeing applied concepts. Also, having taken the AP exam, it was a lot more beneficial to have actually done processes than to have just memorized terms.” However, students reported struggling with several aspects of the project, including the amount of time required to perform the evolution experiments (Module 1) and the sequence analysis activities (Module 2). We identified four themes in the positive and negative responses described below. These responses helped us refine the modules to better feature the aspects students enjoyed and learned the most from and improve the presentation of concepts the students found overly confusing.

### Agency in running experiments

Most students stated that the process of transferring their yeast to fresh media each class period and watching their yeast grow was a positive experience (11/17 surveyed). For example, one student wrote, “It was fun to do the same thing every day and see our yeast grow and get stronger.” They also enjoyed choosing the dose of clotrimazole to which they exposed their yeast, though sometimes this was frustrating when they increased the dose to a level that prevented their yeast from growing and had to go back to an earlier transfer to recover their experiment. These instances provided an opportunity to discuss the limits of their yeast’s drug resistance and reflect on how that changed over the course of the Module 1 experiment. Four of the surveyed students expressed that they appreciated mastering the technique of sterile transfers, saying things like, “It became a routine that we got good at, and we were able to practice scientific procedures.” These findings led us to emphasize student agency in the framing of Module 1 in subsequent implementations.

### Motivated by competition

Module 4 consists of paired competition experiments to find which evolved strain could grow the best in a high drug concentration. The promise of determining which group’s yeast reached the highest fitness in clotrimazole media was a significant motivator for many students (14/17 surveyed), though 3/17 mentioned a dislike of losing. One student wrote, “This part was really fun. I loved competing with the [other] group. It got me really invested in the well-being of my yeast.” As above, we emphasized this activity (Module 4) in subsequent implementations and encouraged students to explore different strategies for maximizing the fitness of their strains.

### Scaffolding of competition experiments

Some aspects of the competition experiments, however, were confusing to students, such as the process of counting colonies (2/17). One route to clotrimazole resistance involves loss of cellular respiration (by mutations in the mitochondrial genome). Respiratory deficiency results in a slow-growth phenotype on media without clotrimazole, a “petite” mutant. These petite colonies were much smaller than their competitors, which was difficult for many students to reconcile with their conceptions of improved fitness in the context of clotrimazole resistance. To address students’ confusion in later classes, we introduced the petite phenotype explicitly through the addition of Module 5, which uses this phenotype as an example of the phenomenon of an evolutionary trade-off: when adapting to one condition leads to decreased fitness in an alternate condition. Additionally, we emphasized that the number of colonies is more important than the size of the colonies. This is because the number of colonies represents the number of live cells in the culture, which corresponds to the success of a culture in producing progeny in the presence of clotrimazole. Colony size only reflects growth rate on agar media lacking drug, which is not the phenotype under selection during the experiment.

### Scaffolding of sequence analysis

One of the most confusing aspects for students was analyzing the DNA sequence data (Module 2); 15/17 mentioned something they disliked about this activity, such as, “It was a little complicated to see some of the mutations” and “It’s a lot of letters and it’s sort of dizzying.” In some cases, the confusion reflected doing authentic scientific research, such as, “Researching the causes of the effects was difficult because there was no real guide as this is a sort of novel experiment so it was hard to find substantial information.” Some of this confusion likely stemmed from using the “raw” format that standard sequence analysis software returns to users. These mutation files included extraneous information, such as a “quality score” that reflects a statistical measurement of whether a mutation call might be a false positive. We also experienced difficulties implementing a sequence analysis and visualization program, the Integrative Genomics Viewer (IGV), such as long loading times on student laptops. These issues distracted from the goal of Module 2. One student described “having to look at the confusing yeast sequence in the confusing program that did not seem to work on anyone’s computer.” In year 2, we simplified the mutation files by removing much of the unnecessary information and focused attention on the literature search component. Students now proceed from a mutant gene name to the *Saccharomyces* Genome Database (yeastgenome.org) to begin exploring gene function (**Supplemental Text 3**).

### Iteration and outcomes from California implementation

These initial modules were revised based on student feedback in 2017-2018. During the 2018-2019 school year, California students participated in Modules 1-5, adding activities about measuring resistance to multiple drug concentrations and evolutionary trade-offs. We collected limited pre-post survey data during this school year (**Figure 3**), which informed continued iterative improvements of the modules, but we do not report them here.

To determine how this intensive activity impacted overall learning in the course, we aggregated AP Biology test scores from AP Biology students at the California school from 2014 to 2019: four years prior to implementation of yEvo and through two years of implementation (**Supplemental Figure 1**). Student scores showed a general upward trend throughout this period, possibly reflecting Liam’s growing comfort with teaching the material to this population. Scores from the years after implementation continued this general trend and produced the highest scores yet. This suggests that the significant time allocation for yEvo-related activities did not detract from student achievement on this standardized test and may have increased proficiency.

### Takeaways from Idaho classes 2018-2020

After our initial implementation, we worked with a public high school in Idaho. Students carried out a modified version of Module 1 (experimental evolution) for 15 weeks during the 2018-2020 school years, with transfers reduced to once a week (**Methods**). Most students additionally completed Module 4 (competitions), though this was interrupted by COVID pandemic-related shutdowns in 2020 (**Figure 3**). In the 2019 to 2020 school year, we surveyed 63 students in three classrooms (two teachers) before and after completion of yEvo modules, of which 41 responded to both the pre- and post-survey. When asked about things they liked, responses generally overlapped with those from Liam’s class, including references to competition and seeing a change over time (e.g., “I like seeing our yeast evolve to resist, and when we looked at it a week later either being so excited seeing the blue color or being disappointed in seeing the yeast dead”). A majority of students reported that they would be willing to do the activity again (∼78%) and that it was fun (∼93%) when asked in the post-lab survey (**Table 10**). When asked about things they did not like about Module 1, the most common theme in responses (10 students) related to the amount of time required, waiting between transfers, or that the process was slow.

### Patterns in term usage pre-post

Our survey included four open-ended questions (Q1-Q4) about genetics and evolution concepts related to yeast evolution. Across all student responses, we noticed an increase in the use of several themes related to more precise terminology from the pre-to post-survey (**Table 4, Table 5, Table 6, Table 7; Figure 4**). For example, on Q1 (*What is a gene?*), we saw a significant increase in the term “code.” Interestingly, more students in the post-survey referred to genes as having a “code” rather than using more vague terminology that we grouped under the theme “make” (**Table 4; Figure 4**), whereas, in the pre-lab survey, students were more likely to use “make” than “code.” This may suggest a more precise understanding of gene and DNA structure and function. On Q3 (*How would you describe evolution?*), students completing the post-survey were more likely (when compared to their pre-survey) to reference “slow” and “mutation” and were less likely to include common misconceptions, which we captured under “naive” (**Table 6; Methods**). This supports a more precise understanding of the concept. Interestingly, we measured a decrease in the use of “adapt” on Q3. On Q4 (*What is the role of mutations in evolution?*), we recorded an increase in the usage of “variation.” These patterns will guide further iteration, as described in the discussion below.

**Table 4.** List of codes, percent responses per teacher, and example student response for question 1: What is a gene? * indicate significant difference

| Code | Emily Pre | Emily Post | David Pre | David Post | Example student response |
| --- | --- | --- | --- | --- | --- |
| DNA | 73.7 | 57.9 | 59.4 | 64.5 | "Genes are parts of DNA that create our body composition." [S008] |
| Code | 21.1 | 52.6 | 15.6 | 32.3 | "A component of DNA responsible for determining appearances and bodily structures." [S024] |
| Make | 31.6 | 15.8 | 40.6 | 22.6 | "A characteristic that makes up who and what you are." [S019]<br>"A strand of DNA that decides someone's physical traits" [S034] |
| Heredity | 26.3 | 10.5 | 15.6 | 29.0 | "A thing that determines your traits passed on from your parents." [S033] |
| Trait | 36.8 | 47.4 | 25.0 | 32.3 | "A gene is a strand of DNA which contains information which decides certain characteristics." [S037]<br>"Genes are made up of DNA. A gene is a physical unit of traits." [S053] |

**Table 5.** List of codes, percent responses per teacher, and example student response for question 2: What is a mutation?

| Code | Emily - Pre | Emily - Post | David - Pre | David - Post | Example student response |
| --- | --- | --- | --- | --- | --- |
| Inherited | 5.3 | 0.0 | 6.3 | 12.9 | "A heritable adaption of a species." [S001] |
| Phenotypic | 15.8 | 21.1 | 9.4 | 3.2 | "A mutation is something that is like unusual to happen like blue eyes" [S002]<br>"Something gets screwed up in the DNA and something weird happens like pinkies." [S015] |
| Genotypic | 100.0 | 78.9 | 84.4 | 100.0 | "A mutation is when one of the Bases pairs up with the wrong base pair. It causes mutation because the two bases don't fit together." [S011]<br>"Something gets screwed up in the DNA and something weird happens like pinkies." [S015] |
| Variation | 5.3 | 0.0 | 9.4 | 6.5 | "A genetic variation" [S041]<br>"...resulting in a variant form..." [S052] |
| Evolution | 10.5 | 0.0 | 0.0 | 0.0 | "A change in the genes of an organism, causing the organism to function slightly differently. These mutations add up over time to create the evolutionary process." [S036] |

**Table 6.** List of codes, percent responses per teacher, and example student response for question 3: How would you describe evolution?

| Term | Emily - Pre | Emily - Post | David - Pre | David - Post | Example |
| --- | --- | --- | --- | --- | --- |
| Adapt | 36.8 | 21.1 | 50.0 | 12.9 | "A slow process of adapting traits needed to survive in an environment" [S006] |
| Vague adapt | 26.3 | 31.6 | 12.5 | 25.8 | "Adopting physical changes to help live in changing environment." [S008] |
| Selection | 26.3 | 5.3 | 12.5 | 12.9 | "The process of natural selection using mutations to create a beneficial output." [S028] |
| Survive | 21.1 | 21.1 | 12.5 | 9.7 | "A slow change in a species over time to optimize the ability to survive and reproduce." [S029] |
| Heredity | 0.0 | 15.8 | 6.3 | 12.9 | "Mutations that sometimes stick and get passed on." [S020] |
| Slow | 31.6 | 52.6 | 37.5 | 61.3 | "A species slowly changing generation after generation to adapt to it's environment." [S025] |
| Mutation | 21.1 | 36.8 | 12.5 | 32.3 | "a series of mutations over many generations" [S042] |
| Species | 36.8 | 31.6 | 40.6 | 51.6 | "evolution is the change of a species at a population level over time." [S004] |
| Naive | 21.1 | 0.0 | 9.4 | 3.2 | "a group of animals that are changing themselves according to their weather, geography, etc." [S027] |

**Table 7.** List of codes, percent responses per teacher, and example student response for question 4: What role do mutations play in evolution?

| Term | Emily - Pre | Emily - Post | David - Pre | David - Post | Example |
| --- | --- | --- | --- | --- | --- |
| Change | 52.6 | 42.1 | 37.5 | 45.2 | "They control how beings change" [S003] |
| Trait | 10.5 | 26.3 | 15.6 | 19.4 | "They make certain individuals more resistant or not resistant to things." [S009] |
| Heritable | 21.1 | 21.1 | 15.6 | 16.1 | "Mutations can be passed into offspring causing population wide evolution" [S011] |
| Selection | 31.6 | 21.1 | 31.3 | 25.8 | "The mutation either help the organism survive and reproduce more or it kills it off because it's unable to survive." [S007] |
| Essential for evolution | 47.4 | 42.1 | 34.4 | 29.0 | "It is what causes evolution, a species can't evolve without some type of mutation." [S013] |
| DNA | 5.3 | 10.5 | 3.1 | 3.2 | "Mutation's role in evolution is, mutations change the DNA of the animal which changes the animal, and these changes help with evolution." [S050] |
| Adaptation | 5.3 | 15.8 | 9.4 | 3.2 | "Mutations create useful adaptations that help animals in their environment" [S005] |
| Variation | 5.3 | 26.3 | 9.4 | 19.4 | "mutations in a populations can lead to change so if there is a species of beetles living on black gravel it starts out with **75 percent white and 25 percent black** beetles the white beetles are more likely to be caught so then the black beetles have more babies and over time the white beetles will die out. "[S004] |

**Figure 4.**
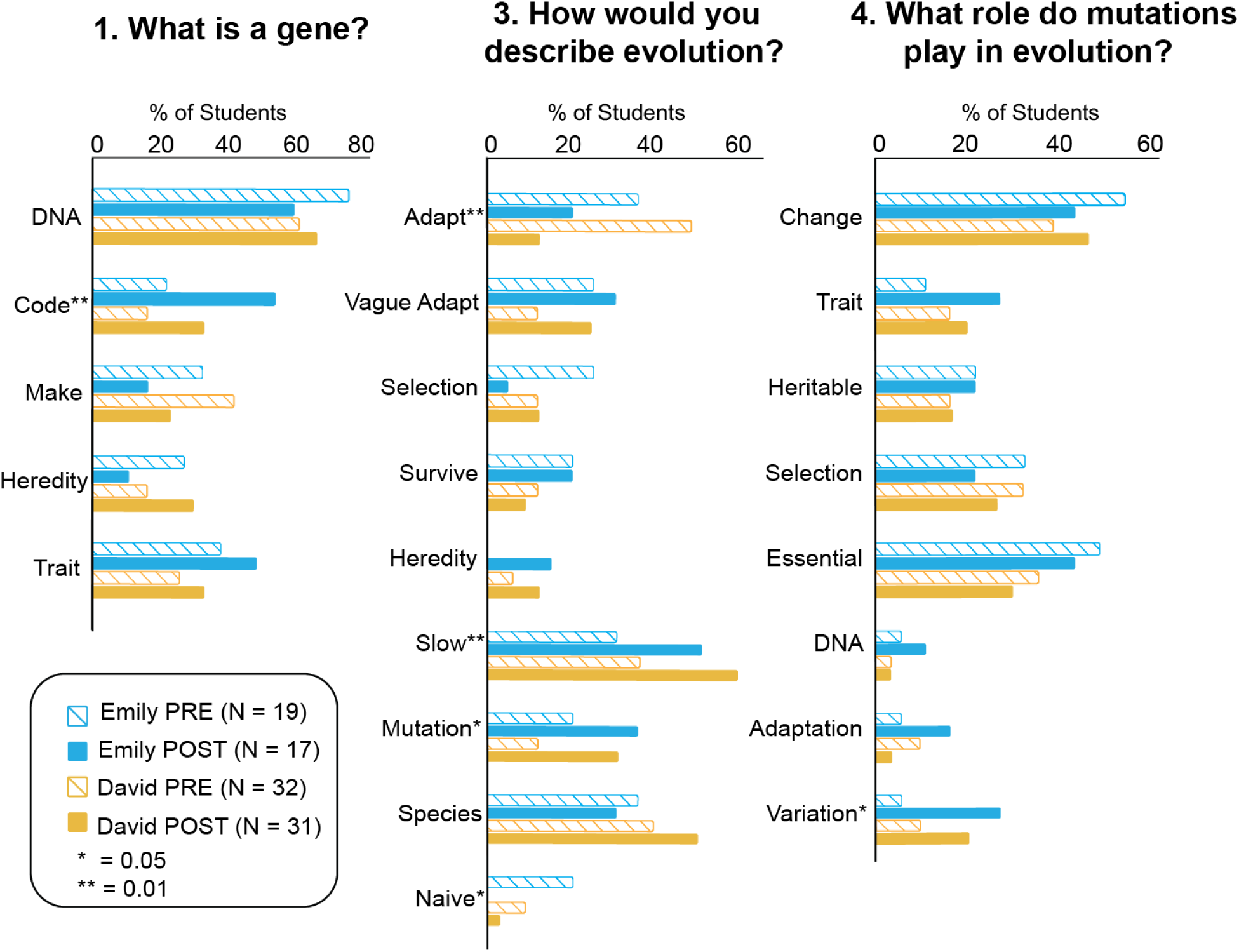
The proportion of the Idaho school students who used key terms and phrases (codes) in their responses to three pre and post-survey questions, grouped by each teacher (Emily, David). Percentages per code do not sum to 100% because a single student could have multiple codes per response. Significant differences between pre and post across all students are indicated by * (alpha 0.05 level) and ** (alpha 0.01 level). See p-values in **Supplemental Table 7**.

### Increase in proficiency in an activity-specific question

Q5 asked students to describe antibiotic resistance. We found that the average score across both teachers’ students of these responses increased by ∼55% after participation in yEvo (0.68 out of 3; p < 0.0001 on a 2-tailed paired t-test; **Supplemental Figure 2**). Though this question asked about antibiotics (general terminology for a drug that is usually associated with bacteria) instead of antifungals, the concepts are nearly identical, suggesting that the students grasped this topic at a conceptual level.

### Response patterns about the mutation process match findings from a previous study

Our Q6 ([*explain why or why not] individual microbes develop mutations in order to become resistant to an antibiotic and survive*) was modeled from Richard et al. 2017, which used it to assess reasoning on mutation processes in evolution in science students and professors. It asked students if they agree (Likert scale) that individual microbes mutate to become resistant to an antibiotic and then asked them to justify their responses, which was coded similarly to the scheme used in Richard 2017 (**Methods**). After completing yEvo, more students disagreed with the statement (12/41 pre vs. 19/41 post; **Supplemental Figure 3**). When comparing post-lab to pre-lab responses, we saw an overall increase in the use of the codes “individual,” “random,” and “natural selection,” and a decrease in “purpose” (**Table 8**; **Supplemental Figure 3**). Students who agreed with the statement were more likely to describe the mutation as serving a purpose (**Supplemental Figure 3**).

**Table 8.** List of scores, percent responses per teacher, and example student response for question 6: [Explain why or why not] individual microbes develop mutations in order to become resistant to an antibiotic and survive.

| Term | Emily - Pre | Emily - Post | David - Pre | David - Post | Liam - Pre | Liam - Post | Example |
| --- | --- | --- | --- | --- | --- | --- | --- |
| Individual vs. Population (distinguish) | 10.5 | 47.4 | 31.3 | 45.2 | 53.8 | 50.0 | "not individual microbes but colonies" [PP022]<br>"Individual microbes can develop mutations, but those mutations may benefit or hurt them. If it benefits them then with will help the next generation become resistant, but if it hurts them, they will die and it doesn't help any generations." [PP044] |
| Random | 0.0 | 15.8 | 18.8 | 51.6 | 61.5 | 33.3 | "They don't develop mutations, but it happens randomly." [PP005]<br>"First of all, mutations aren't intentionally done." [PP018] |
| Purposeful | 42.1 | 26.3 | 37.5 | 19.4 | 15.4 | 33.3 | "Microbes mutate based on the needs of the species, so yes they mutate to be more resistant to antibiotics "[PP099]<br>"they need to mutate in order to fight the antibiotic or else they won't survive." [PP043] |
| Natural selection | 36.8 | 57.9 | 12.5 | 29.0 | 53.8 | 50.0 | "If a mutation happens to make the microorganism more resistant to an antibiotic, then it is more likely to survive and reproduce. Over time, most of the population will become resistant to the antibiotic." |

### Increase in confidence in the ability to design a valid biological experiment

One of our central goals was to put the tools of science in students’ hands. After completing Module 1, 18/46 (39%) students (all collected post-surveys regardless of pre-survey completion) reported increased confidence in their ability to design a valid biological experiment, compared to three who listed a decreased confidence [**Table 9**]. In interviews, students expressed that they appreciated learning how to work with yeast and observing how yeast responded to different environments, including statements like, “It was a big excitement every time we see a growth in our tube.” These observations align well with survey responses from students in the California school regarding an appreciation for their newfound mastery of basic microbiology techniques.

**Table 9.** Changes in likert scale questions between pre- and post-lab surveys for David and Emily’s students combined. We converted a likert scale (strongly disagree - strongly agree) to a numerical 1-5 scale and calculated the difference between paired pre- and post-lab responses for each student who completed both (N = 41).

| Difference | I am interested in a career in biology. | I am interested in a career in STEM. | I am confident that I can design a valid biology experiment. | I think about the biology I experience in everyday life. | The study of biology is only useful when it directly benefits human health or well-being. | Biologists may make different interpretations based on the same observations. | Experiments that are done under lab conditions can provide information that applies to the real world. |
| --- | --- | --- | --- | --- | --- | --- | --- |
| -3 | 0 | 0 | 0 | 1 | 0 | 0 | 0 |
| -2 | 1 | 0 | 2 | 0 | 3 | 0 | 1 |
| -1 | 8 | 7 | 1 | 9 | 5 | 4 | 8 |
| 0 | 22 | 23 | 23 | 20 | 25 | 27 | 22 |
| 1 | 8 | 9 | 10 | 6 | 6 | 8 | 6 |
| 2 | 2 | 1 | 2 | 3 | 0 | 2 | 3 |
| 3 | 0 | 0 | 3 | 3 | 2 | 0 | 1 |
| 4 | 0 | 1 | 0 | 0 | 0 | 0 | 0 |
| Mean difference | 0.05 | 0.20 | 0.44 | 0.30 | 0.02 | 0.20 | 0.12 |
| Pre-lab mean | 2.85 | 3.61 | 3.29 | 2.93 | 1.90 | 4.32 | 4.00 |
| Post-lab mean | 2.90 | 3.80 | 3.73 | 3.23 | 1.93 | 4.51 | 4.12 |

### Lasting impact on interests in STEM fields

Since these activities were designed to be true to scientific research practices, we investigated how participation would impact student conceptions of the process of biology (**Supplemental Figure 4**) and interest in STEM fields (**Supplemental Figure 5**). On pre- and post-lab surveys, we asked students about their interest in biology or STEM careers. Students who completed both surveys showed a small average increase in interest over this time period [**Table 9**], though this difference was not significant. We directly asked students whether participating in this activity increased their interest in a STEM or a Biology career on our post-lab surveys. Of students who responded to both the pre- and post-survey, 15/46 (33%) agreed or strongly agreed with one or both questions [**Table 10**]. This encouraging result led us to wonder what students would recall one year later.

**Table 10.**
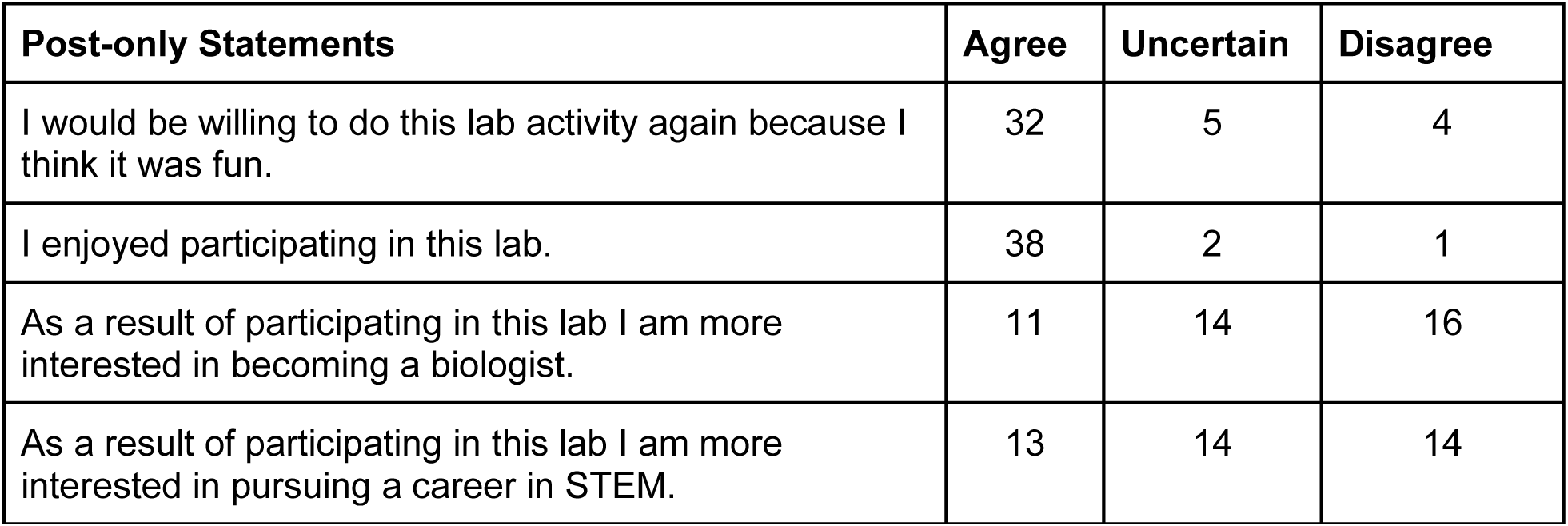
Number of students who responded as agree or strongly agree (“agree”), uncertain, or disagree or strongly disagree (“disagree”) for each of four statements asked in the Idaho post-survey (N = 41).

| **Post-only Statements** | **Agree** | **Uncertain** | **Disagree** |
| --- | --- | --- | --- |
| I would be willing to do this lab activity again because I think it was fun. | 32 | 5 | 4 |
| I enjoyed participating in this lab. | 38 | 2 | 1 |
| As a result of participating in this lab I am more interested in becoming a biologist. | 11 | 14 | 16 |
| As a result of participating in this lab I am more interested in pursuing a career in STEM. | 13 | 14 | 14 |

### Feedback from students one to two years later

Thirty-nine students from the 2018-2019 and 2019-2020 school years at the Idaho school were surveyed again in 2021 (**Supplemental Table 8**) to check for retention of knowledge gained from the yEvo experiments. When asked about their memory of the lab, students mentioned: “yeast poison” or “acid” to describe the clotrimazole, showing that the students seemed to retain the information of the general experiment but not the specifics of the lab. The majority of students (27/39) responded that the lab was valuable to their experience. Four students felt that their experience lacked an understanding of how the research stemming from their experiments was being used. Two students mentioned losing interest or motivation to continue the repetitive task of transferring yeast over the course of the lab, especially if they were not pushed to continue discussing it. However, ten students mentioned how it demonstrated applications of evolution or simulated real lab experiences: “It mimicked actual lab work, and we got to make real observations (we actually got to observe yeast evolving).”

### Idaho Teacher feedback

After completing the 2019 to 2020 school year, we interviewed the two teachers from the Idaho school about their use of the lab activity. Both teachers had assistance from University of Idaho researchers. They felt that both the lab activity and increased access to trained microbiologists assisted them in teaching aspects of biology that complemented their expertise. As in the student responses, the teachers noted that the competition was a driving factor for student engagement. The teachers liked that the lab gave students freedom in their choices at various steps and allowed students to see evolutionary changes happening quickly. As students gained proficiency with the sterile transferring protocols, they carried out daily or weekly tasks with little assistance in approximately 10-15 minutes. As a result, these teachers reported that incorporating the lab did not overburden their teaching.

Suggestions from these teachers included developing a resource to help the teachers understand the biology of yeasts to answer student questions better. The teachers noted that online videos or resources might increase the ease of running the lab, particularly if access to university researchers was limited in future implementations. They also mentioned challenges with data collection and requested a standardized data log to improve ease of management and organization once the samples were sent back to the university to be processed. Teachers also showed interest in the students having more control in the design of their experiments (e.g., choosing a selection pressure) to add additional agency. The protocols provided with this manuscript (**Supplemental Texts 1-6**) reflect these requests.

## Discussion

### Alignment of yEvo goals and outcomes

Here we outlined the development, implementation, and preliminary evaluation of curricular modules that engage high school students in authentic experimental evolution research. Our design process was greatly aided by close collaborations between experienced teachers and university research labs. This collaboration allowed us to formally and informally monitor how the yEvo lab curriculum impacted student motivation and achievement. We were then able to adjust our approach based on feedback from students and teachers.

We had two primary concerns about initiating yEvo in high schools that motivated our evaluation program. First, **does the amount of time required for these exercises detract from other educational goals?** The California teacher (Liam) co-developed Module 1. He carried it through the entire school year from 2017 to 2020 and estimated that it took up approximately 20% of total class time. Despite this, participation did not harm standardized test scores (**Supplemental Figure 1**). This suggests that the benefits of an authentic research project compensated for a reduction in time for traditional content and that teaching some of this content via yEvo was successful. More investigation is needed to determine if this holds in multiple classroom contexts.

Our evaluations indicate that students at our second school in Idaho, who utilized a streamlined protocol that required less class time, improved their grasp of activity-specific concepts and increased the use of terminology related to several learning objectives. Our findings from Q6 (*[explain why or why not] individual microbes develop mutations in order to become resistant to an antibiotic and survive*) are in line with those of Richard 2017, which showed that undergraduates who disagreed with the statement were less likely to display teleological (purposeful) reasoning, a common misconception about evolution. Critically, the use of teleological reasoning decreased after completing Module 1. Students were more likely to disagree with the statement, which the Richard group characterized as an expert-like response. This could explain a curious result on Q3: that students were less likely to use terminology related to adaptation in their descriptions of evolution. This may be because students were more likely to consider evolution as a random process that results in increased fitness rather than a purposeful adaptive process. In conclusion, we found that yEvo, although requiring a significant investment of time by teachers, resulted in a beneficial and authentic research experience for high school students at two different institutions.

Our second major question was: **does the repetitiveness of Module 1 impact technical confidence and interest in biology?** The length of time for which students carry out these experiments may lead to boredom, and some student responses on the post-activity survey indicated this. For example, one said, “I feel like the lab had no set end goal, so it sort of just petered out and we lost interest.” We see this as a worthwhile risk since the repetitive experimental design accurately reflects many research methodologies. Further, the repetitiveness allows students to improve their proficiency in the relevant procedures. Students’ experiences with these methods may help them better evaluate their interest in a science career and gain essential skills for pursuing a science career. Critically, students on average, stated that participation in Module 1 contributed to increased confidence in their ability to design biological experiments and an increased interest in Biology/STEM careers (**Table 9**). Even after one to two years, students could recall the techniques utilized in the lab consistently, though they did not use precise terminology in describing these procedures. Further, a majority (27/39) felt that the lab exercise had been a valuable experience. Each of these points may be related to their opinion that they had mastered the techniques required for Module 1.

In addition to these positive signs of student learning, our educational collaboration was beneficial to the research goals of participating researchers. A large body of research exists on clotrimazole resistance (and azole resistance more broadly) in a variety of species of yeast (Gulshan 2007; Demuyser 2019). This prior work enabled us to evaluate and contextualize student results. Carrying out these investigations with several classes of students provided a high degree of replication and led to our team identifying new genetic factors that contribute to clotrimazole drug resistance in yeast (Taylor 2021). In short, we isolated 99 clones from Module 1 experiments performed by 203 students from the California and Idaho schools over three years. These were subjected to further analysis and whole-genome sequencing. Every student clone we sequenced possessed at least one mutation that fits known resistance mechanisms to clotrimazole (Taylor 2021). This result demonstrated that our protocol could reliably reproduce clinically-relevant findings from traditional research laboratories. In addition, student data suggested previously uncharacterized mechanisms of resistance, providing new insights into clotrimazole resistance (Taylor 2021). As a result, future students participating in yEvo will be able to see a lasting impact on the field.

It can be unappealing for researchers to take time away from academic laboratory research to develop educational activities, but our experience (and many others, e.g. (Kerfeld 2007; Jordan 2014; Brownell 2015; Saha 2017; Mavor 2018)) demonstrates that education and research can be merged in mutually beneficial ways. For instance, the sequencing reactions in Module 2 are the most costly aspect of the entire lab (at least $20/sample), a cost the partner labs covered, since these experiments were relevant to ongoing research. As a result, these data serve multiple purposes: education/training and research.

### Implementation considerations

The modules we present cover advanced topics and methods that can be intimidating for first-time users. To develop them, we worked with teachers possessing extensive teaching experience and training. The teacher at the California school has a doctorate and one of the Idaho teachers a master’s degree, each in a subdiscipline of biology. In interviews after implementing the evolution module, teachers at the Idaho school reported that the close collaboration with researchers, and the presence of an undergraduate student facilitator at the Idaho school, increased their comfort. Though we did not measure it directly, evidence exists that peer facilitators can benefit students (Sellami 2017). We expect that the interaction with researchers makes the lab feel more authentic than standard classroom exercises. From the researcher’s standpoint, the close collaboration gave us more insight into aspects of this project that needed improvement, which we may not have learned without extensive interactions. For those interested in developing similar collaborations, specific suggestions for developing fruitful teacher-researcher collaborations can be found in (Warwick 2020; Knippenberg 2020).

### Future development of yEvo curricula

A critical next step will be to rigorously evaluate the impact of each of the modules we have developed. Since these modules can stand independently, we anticipate that teachers will utilize distinct combinations based on their course objectives, interests, and resources. This would create natural experiments in implementation through which we would be able to tease out the impact of individual modules by comparing learning gains in classes with or without a particular module. Since the Module 1 experimental evolution activity seemed to be polarizing (students reported differences in the aspects that were motivating), it is prudent to investigate how various module combinations or implementation strategies impact motivation.

Future lessons could focus on how researchers are using these data. In early iterations, we collected data that has since played a role in published investigations. Introducing this process could show students how research projects are structured. We could additionally incorporate opportunities for students to develop follow-up experiments or design entirely new experiments based on the experimental evolution paradigm.

The Module 1 experimental evolution framework can be applied to any environmental condition that supports yeast growth. This opens the door to framing experiments around applications of yeast biology such as food or biofuel production. However, not all conditions will be equally valuable in a teaching context. The clotrimazole resistance mutations our students isolated have a strong effect, making it possible to see a difference between evolved and ancestral strains after only a few transfers. The most common mutations in clotrimazole resistance impact processes that are relatively straightforward to contextualize due to decades of research on this topic. Prototyping new conditions in the university lab will be a useful precursor to classroom deployment to identify which conditions are most amenable to both teaching and research goals.

### Impact of evolution research experiences

The basic conceptual and technical skillset we aim to impart will be of value for students regardless of whether they continue in biology. Skills related to Module 1, such as sterile technique, liquid handling, and record-keeping, will be useful in many modern biology applications. Whole-genome sequencing and related genomic technologies covered in Module 2 are being incorporated into all subdisciplines of biology. Concepts such as the effect of sequence variation on traits, the evolution of drug resistance, and the role of molecular diversity on vaccine efficacy are increasingly making their way into the public sphere. Familiarity with these concepts will certainly benefit public health and understanding of genetic results encountered in health care and direct-to-consumer settings.

We believe that the yEvo experience is a window into the process of evolution that can reinforce concepts throughout high school biology curricula. We designed these modules to illustrate that evolution is an ongoing process that we can study and measure using model organisms with a short generation time. This may open up a route for discussion on evolution with students skeptical of standard evolutionary models. The agency of working with one’s own case study in evolution may provide motivation and scaffolding for future learning. Demonstrating how experimental tools are used in modern biology labs may inspire deeper thought into what is currently possible in biological research. These hypotheses will require investigation, and our yEvo framework provides a valuable tool in this endeavor.

## Acknowledgments

We thank Jef Boeke and Jasmine Temple for color plasmids, Sayeh Gorjifard for figure design assistance, Naomi G. Moresi for the experiment photographed in Figure 1D, and all participating schools and teachers.

## IRB number

STUDY00003148

