## Supplemental materials for "yEvo: a modular eukaryotic genetics and evolution research experience for high school students"

**Supplemental Text 1.** Curriculum overview.

**Motivation**

The goal of yEvo is to involve students in an authentic research experience that connects genetics, cellular biology, and organism-level phenotypes in an evolutionary context. Briefly, students select for yeast mutants that are more resistant to a "stressor" (any environmental condition that slows growth), examine sequencing data to identify mutations that may be responsible for this resistance phenotype, and use cellular/molecular models to contextualize how their mutations may be connected to the resistance phenotype. These are complex topics with which (in our experience) students frequently struggle. By focusing on **their** experiments and **their** mutations, we expect this experience will provide scaffolding and motivation for further learning.

Experimental evolution can be utilized to study adaptation to any environmental condition that supports the growth of yeast. In our first implementations, we utilized an azole-class antifungal called clotrimazole. Azoles are one of the most commonly used antifungals in medicine and agriculture, and azole resistance is a growing global health crisis. A large body of research exists on azole resistance in a variety of species of yeast, so we and our students can find published information about their mutations to provide context to their results. At the same time, few studies have applied experimental evolution to identify genetic factors contributing to azole resistance, so we felt there would be opportunities for our students to make novel discoveries. We encourage you to consider other experimental evolution conditions and would be happy to support your efforts!

**Module 1: Evolution.** As the ancestral populations for our evolution experiments, we used a collection of yeast strains that were previously engineered to express vibrant pigments by members of the Boeke lab at New York University. This allowed us to watch for contamination of growing cultures by monitoring the color, and was also used for the competition assays in module 4. We added a secondary antibiotic (G418) to reduce the odds of a contamination event and to maintain the plasmids on which the pigment genes are carried.

Students first grew yeast in the presence of an over-the-counter antifungal azole drug (FungiCure) for several weeks, performing transfers into fresh drug-containing medium at regular intervals. As students observed improved growth, they increased the drug dosage. By the end of the evolution experiment, students' evolved yeast typically grow more robustly in the presence of the antifungal drug due to mutations that increased their level of resistance. Evolved yeast typically survived exposure to much higher concentrations of the antifungal drug (4-16x) than their unevolved ancestors. This result provided an opportunity for qualitative ("In dose X, the evolved culture is 'cloudier' than ancestral strain") or binary ("Both strains grow in dose X but only the evolved strain grows in dose Y") comparisons. The length and frequency of interaction with these experiments is flexible to classroom time constraints, as resistance phenotypes can be reliably observed after 5 transfers, which can be performed at intervals of 2 days or up to 2 weeks.

Students carried out evolution experiments for 7 to 34 weeks depending on the year and the classroom using one of two protocols. We were able to isolate clones with increased azole resistance from some experiments as early as two weeks.

**Safety.** Baker's yeast are generally considered nonpathogenic, but isolates have been obtained from hospital patients suffering from complications of compromised immune systems. Drug resistance and pathogenicity are distinct traits, and the laboratory strain we work with in yEvo lacks several characteristics that clinical isolates or pathogenic species possess, such as ability to form biofilms and to grow robustly at human body temperature. Still, it is essential that care be taken in this exercise, particularly around sterilization and disposal of materials.

To prevent contamination by foreign microbes, we recommend utilizing media that can prevent growth of organisms other than the lab strain of *S. cerevisiae*. Our experiments have included a drug called G418, which our particular laboratory strain is resistant to. No method of preventing contamination is perfect, so it is critical to closely observe cultures for signs of contamination. Yeast cells will settle to the bottom of a test tube after 20 minutes in a stationary test tube rack, forming a pellet. Signs of contamination include if the color of the pellet changes, or if cells do not completely settle.

**Module 2: Genomics.** In our trials, teachers sent their evolved yeast to the University of Washington, where we used whole-genome sequencing to identify mutations that may play a role in adaptation. The types of DNA changes represented by these mutations also allowed exploration of the mutational process. Students recovered strains with single base changes, small deletions and insertions, new transposon insertions, and DNA copy number changes ranging from small segments to entire chromosomes. The evolved strains contained mutations in both the nuclear and mitochondrial genome, emphasizing aspects of cell biology present elsewhere in the curriculum.

Students were provided with information about mutations present in evolved yeast that occurred during their or another class's evolution experiments. They performed a collaborative literature search on mutated genes, and on antifungal resistance broadly, and built hypotheses about how these mutations may impact stress tolerance. This search was aided by the *Saccharomyces* Genome Database (SGD; yeastgenome.org), which curates data and publications from decades of research in yeast. Students used a molecular model of azole drug resistance we have developed to contextualize their mutations.

Yeast can become resistant to azoles through mutations in many cellular processes, but two mechanisms dominate both student experimental results and sequencing of clinical isolates of pathogenic species of yeast. One is through mutations that increase the amount of Erg11 present and thus offset the inhibitory effects of the drug. The second is through mutations that increase expression of membrane proteins called drug efflux pumps that can remove azoles from the intracellular environment. Students were consistently able to identify connections between their mutations and these known mechanisms.

**Modules 3 and 4: Fitness.** The next two modules focused on yeast evolutionary fitness. In the minimum inhibitory concentration (MIC) module, students inoculated their evolved yeast and ancestor into media containing several concentrations of antifungal drug. After 2-7 days, students examined each culture to determine the minimum drug concentration that inhibited growth of each strain. The evolved strain was expected to grow more robustly (as observed by the size of yeast "pellet" at the bottom of a test tube or cloudiness of culture after shaking) than the ancestor in higher azole concentrations.

In the competitive fitness module, students used competition experiments to assess relative fitness of independently evolved strains. Evolved strains of different colors were mixed and grown together in a liquid growth medium with and without azole. The cultures were then diluted and plated onto agar media plates. Each individual cell on this plate will form a colony that expresses each strain's distinctive color, enabling counting of colonies specific to each strain. A strain that produces more colonies can be said to have a higher competitive fitness than the strain it was mixed with, because more cells from that strain were present in the mixed culture at the end of a period of competitive growth. To frame the activity, we tasked students with identifying the most resistant evolved populations. Student groups were paired in a tournament-style bracket in which the "winner" of each competition moved on to compete their yeast with another winning group.

**Module 5: Fitness Tradeoffs.** Finally, students used a phenotype linked to resistance to visualize a tradeoff in fitness. Evolved yeast have increased tolerance to the antifungal they were selected in, but this antifungal resistance can come at the expense of other traits, leading to decreased ability to grow in alternate environments. In the most extreme example, many antifungal-resistant clones were unable to grow on media in which glycerol is their primary carbon source. Students grew their yeast on an agar plate with dextrose as the carbon source (permissive), picked colonies of yeast, and transferred them to a plate with glycerol as the carbon source (selective). Students recorded the percentage of randomly-chosen colonies that were unable to grow on glycerol media, indicative of a tradeoff between the ability to grow in the presence of an antifungal and the ability to utilize glycerol. The frequency of this phenotype varied widely across experiments and timepoints, another sign of population-level change due to natural selection of new mutations.

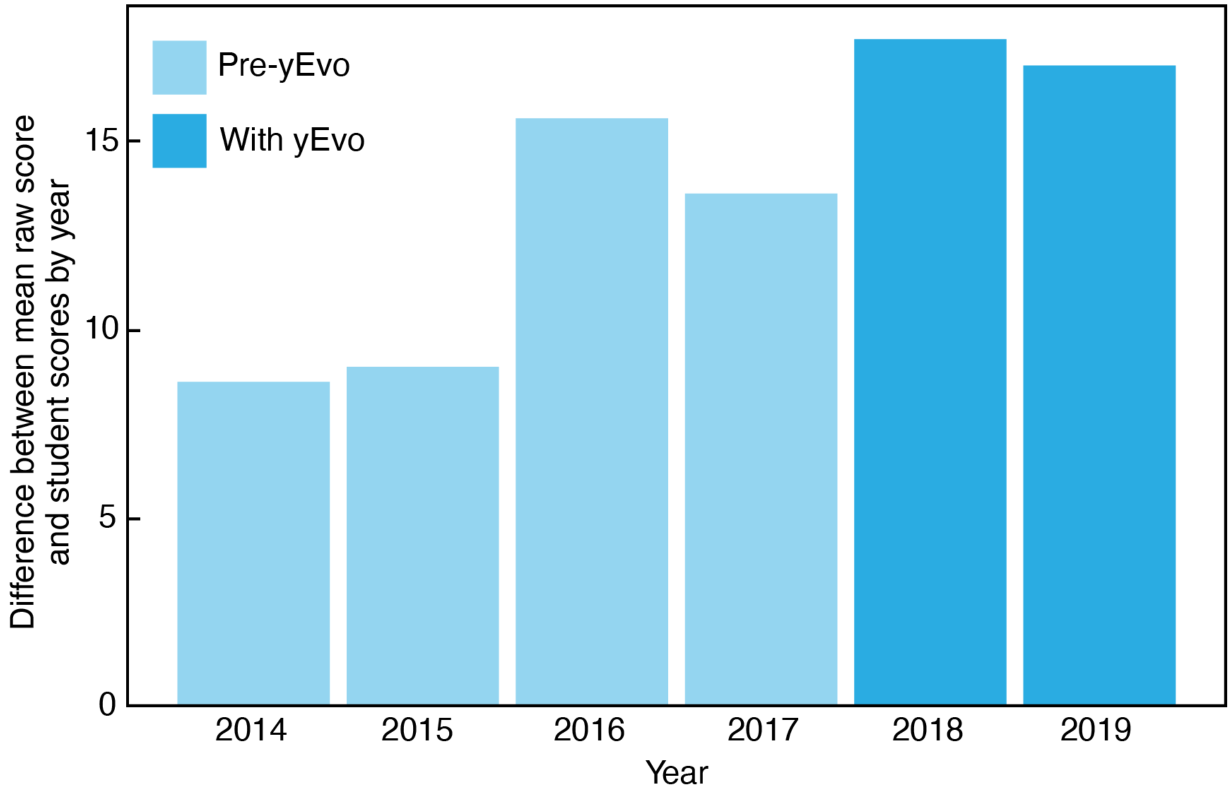

**Supplemental Figure 1.** AP scores of California school students over six years.

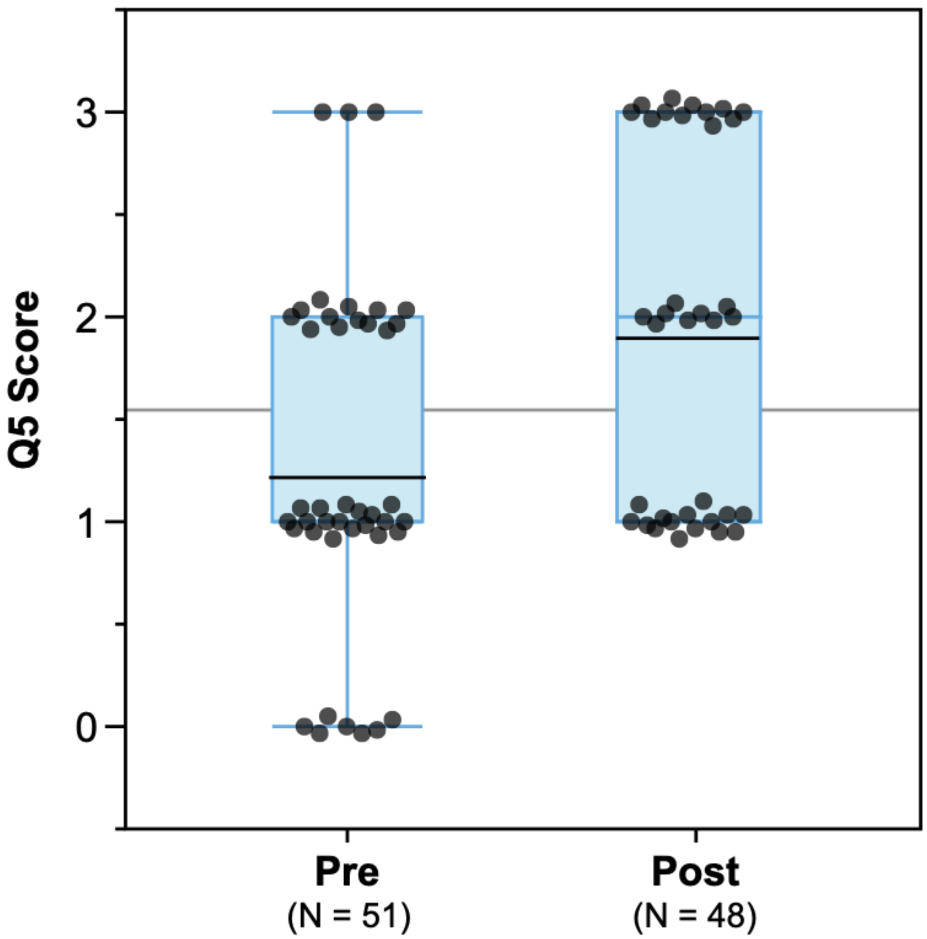

**Supplemental Figure 2.** Box plots of question 5 (Q5) scores for the pre (N = 51) and post (N = 48) responses to the question *How would you explain antibiotic resistance to a fellow student in this class?* Responses on the pre- and post-lab survey were given a score of 0,1, 2 or 3 based on their correctness. Points have been distributed vertically and horizontally to reflect the density of responses given a particular score. Average scores for pre and post indicated by a black line; global average gray middle line.

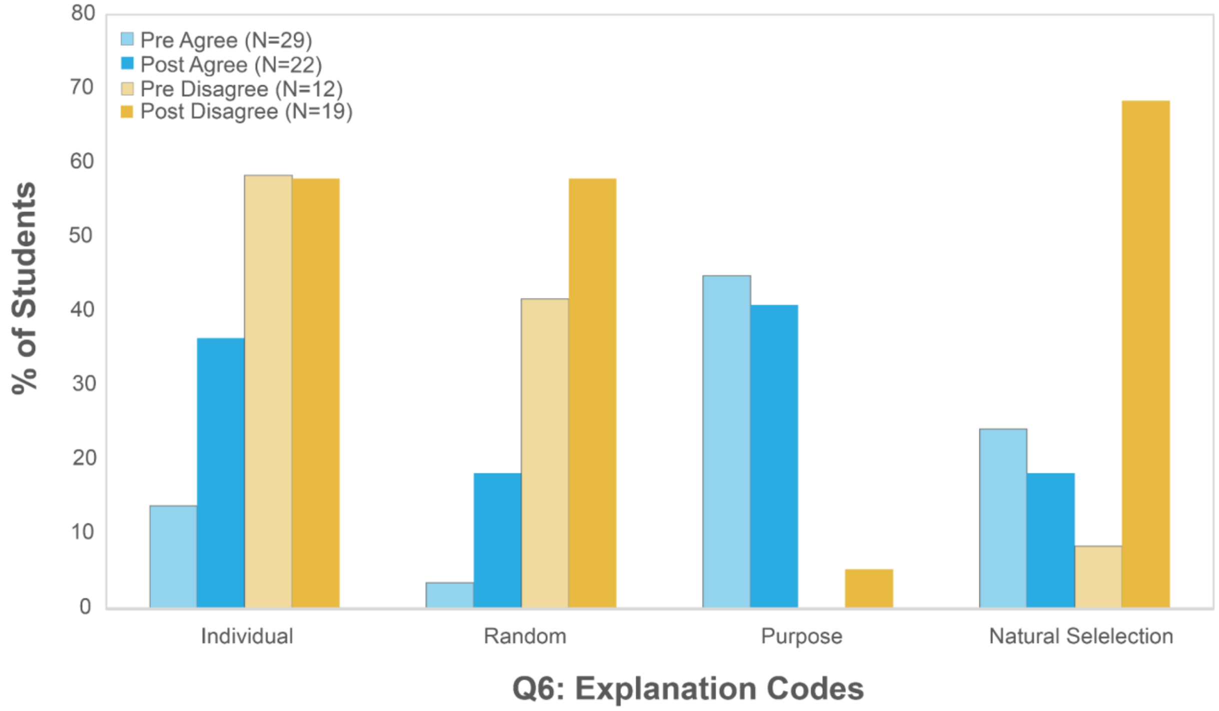

**Supplemental Figure 3.** Percent of student open-ended responses to question 6 (Q6; [*Explain why or why not] microbes develop mutations in order to become resistant to an antibiotic and survive*.) coded by four explanations (see **Table S6**) for pre- (lighter shade) and post- (darker shade) surveys. Each student's response (N = 41) is categorized by their level of agreement to the original statement; agree (blue; either agree or strongly agree) and disagree (yellow; either disagree or strongly disagree).

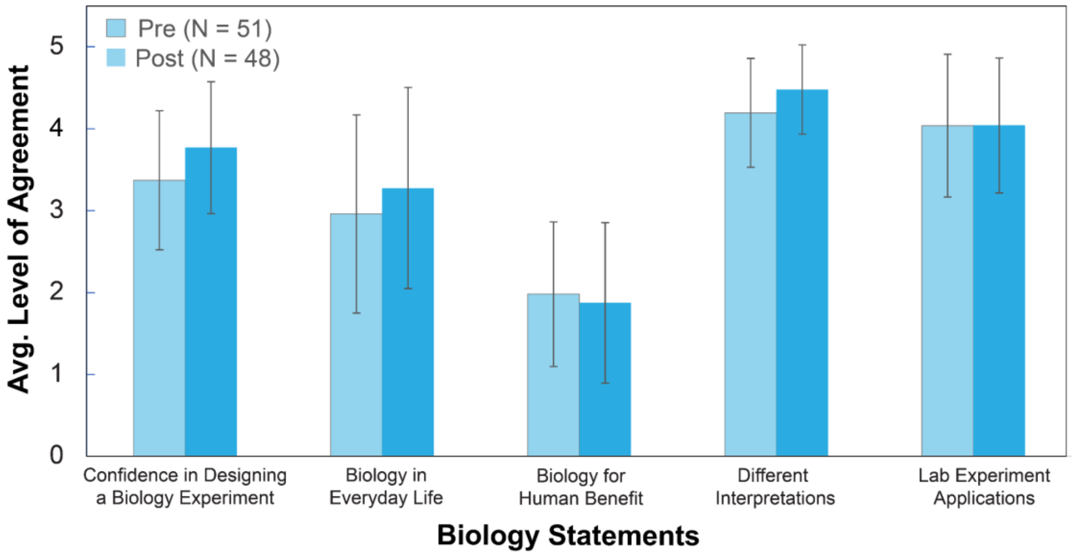

**Supplemental Figure 4.** Average level of agreement (5 = strongly agree) to Likert-response biology statements from students who completed a pre- (N = 51) or a post-lab (N = 48) survey at the Idaho school during the 2019-2020 school year. Standard deviation shown. None of the statements were significantly different from pre to post (t-test, alpha = 0.01).

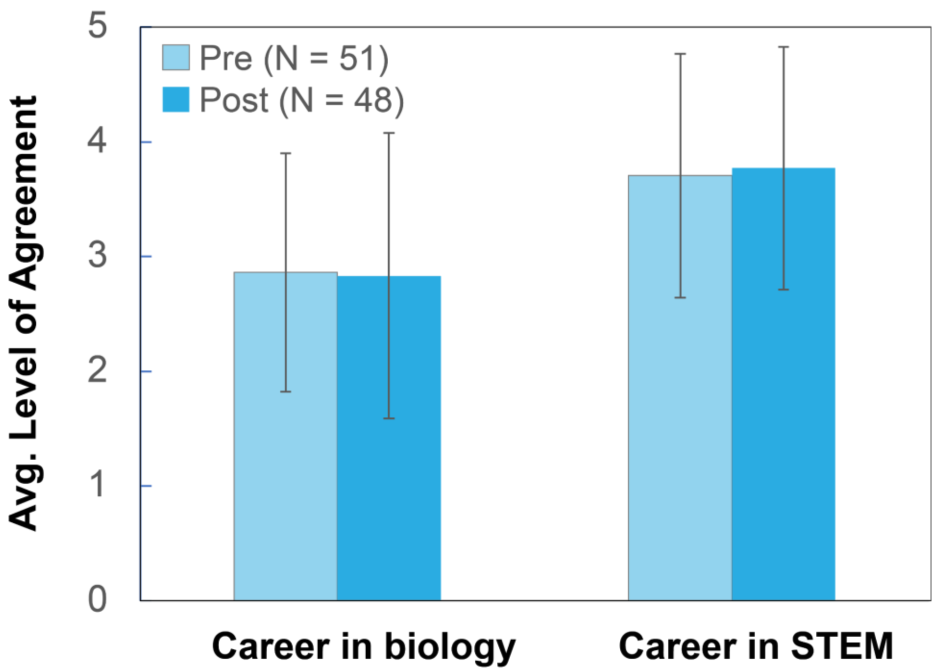

**Supplemental Figure 5.** Average level of agreement (5 = strongly agree) in Idaho school students' interest in either a career in biology or a career in science, technology, engineering or math (STEM) in the pre- (N = 51) or a post-lab (N = 48) survey at during the 2019-2020 school year. Standard deviation shown. Neither of the statements were significantly different from pre to post (t-test, alpha = 0.01).

| Name of code | Description |
|---|---|
| DNA | Described gene as made up of DNA, is DNA, or is a section / sequence of DNA. |
| Code | Mentioned that a gene contains information or codes for / determines the traits of an organism. Other verbs allowed including ‘decides’ / ‘determines’ / ‘controls’ traits.<br>Did not allow ‘associated’. |
| Heredity | Mentioned genes are the unit of heredity or that they are inherited or come from / are passed on by parents. Did not have to use the term heredity or inherited. |
| Trait | Referenced that a gene is a trait or that genes determine traits. Allowed reference to a trait, characteristic, feature, structure, or appearance of an organism. It was not sufficient to say that a gene determines ‘something’ or that it ‘makes you who you are’. Must be more specific. |
| Make | Used the word ‘make’ or other word / phrase that skipped over the process steps (intermediate steps) between gene and trait. Allowed ‘create’. Did not allow ‘information’, as this falls under “code”. Did not allow that a gene is ‘made’ of DNA or ‘genetic makeup’. [S023, S027]. |

**Supplemental Table 1.** List of code descriptions for question 1: What is a gene?

| Code | Description | Example student response |
| --- | --- | --- |
| Inherited | Used term(s) such as 'inherited', 'passed down', or 'heritable' when referring to mutations or traits. Allowed responses that implied heritability. | "A heritable adaption of a species." [S001] |
| Phenotypic | Mentioned a change in phenotype or trait. Included references to traits being physically expressed in a different way. Must reference a physical change. | "A mutation is something that is like unusual to happen like blue eyes" [S002]<br>"Something gets screwed up in the DNA and something weird happens like pinkies." [S015] |
| Genotypic | Mentioned any change in genotype. Allowed differences in DNA base pair or nucleotide sequence or mistake / error in copying or replicating DNA. | "A mutation is when one of the Bases pairs up with the wrong base pair. It causes mutation because the two bases don't fit together." [S011]<br>"Something gets screwed up in the DNA and something weird happens like pinkies." [S015] |
| Variation | Mentioned variation among individuals. Allowed use of the phrase 'genetic variation' or description of a mutation as something which creates variation. | "A genetic variation" [S041]<br><br>"...resulting in a variant form…" [S052] |
| Evolution | Used the term mutation to describe how evolution occurs or as something which contributes to evolution | "A change in the genes of an organism, causing the organism to function slightly differently. These mutations add up over time to create the evolutionary process." [S036] |

**Supplemental Table 2.** List of code descriptions and example student responses for question 2: What is a mutation?

| Code | Description |
|---|---|
| Adapt | Mentioned terms ‘adapt’, ‘adaptation’, or ‘adapted’ to explain evolution. |
| Vague adapt | Mentioned adaptation without explicit use of the term ‘adapt’. For instance: ‘evolve to their environment’ or ‘changed to fit the environment’. |
| Selection | Used the term ‘natural selection’. |
| Survive | Mentioned survival of organisms, usually regarding increased survival as a result of a particular trait. Allowed reference to the opposite of survival - death - as a result of species with undesirable characteristics. Did not allow ‘stronger’ traits. |
| Heredity | Explicitly mentioned the terms/phrases ‘inheritance’, ‘heredity’ or ‘passing on’ traits. Allowed reproduction or passing traits to offspring, even if they did not specifically mention genetic factors like mutations. Similarly, allowed loss of traits over time (‘breeding out’) because organisms with those traits die off (do not reproduce). |
| Slow | Mentioned evolution as slow or gradual changes with phrases like ‘over many generations’, ‘long time’, ’long process of time’, ‘many generations’, or ‘change over time’. Did not allow ‘process of change’ or ‘several years of adaptations’. |
| Mutation | Mentioned source of variation from mutations. Allowed ‘Genetic changes’ or ‘changes to DNA’. |
| Species | Mentioned a change in how frequent or common a trait or mutation is in the **species** or **population**, or that change is not occurring at the individual level. Allowed group of animals (more descriptive than ‘animals’ or beings). |
| Naive | Naive explanations mentioned ‘need’ (traits that are ‘needed’ to survive) or ‘intention’ (traits that are ‘chosen’ to help an organism survive). |

**Supplemental Table 3.** List of code descriptions for question 3: How would you describe evolution?

| Code | Description |
|---|---|
| Change | Mentioned mutations lead to changes in individuals or cause differences in organisms. |
| Trait | Referenced a gene is a trait or that genes determine the code for traits. This particular code is more about the reference to a trait, characteristic, feature, structure, or appearance of an organism. It is not sufficient to say that a gene determines codes for ‘something’ or that it ‘makes you who you are’. Must be more specific. |
| Heritable | Stated that mutations are heritable or can be passed down, so any beneficial or deleterious consequence will apply to offspring as well. |
| Selection | Described how mutations can impact survival or the production of more or less progeny. Did not need to use the term ‘selection.’ Did not allow ‘fit’ without additional clarification of differential survival. |
| Essential for evolution | Described how mutations ‘enable’ evolution. Allowed references that mutations are important but did not require an explanation how. Allowed more general language like ‘help’, ‘allow’, and ‘create’ evolution. |
| DNA | Made a reference to changes in DNA. |
| Adaptation | Used the term ‘adaptation’ or ‘adapt’. |
| Variation | Used the term ‘variation,’ described the change in frequency of a trait within the population, or mentioned differences between individuals / organisms within the population. |

**Supplemental Table 4.** List of code descriptions for question 4: What role do mutations play in evolution?

| Score | Description |
|---|---|
| 0 | Responded that they don’t know, OR the student’s response is entirely / mostly incorrect. |
| 1 | Echoed the question without giving any additional information. An example of this would simply be restating that the bacteria are resistant to the drugs. |
| 2 | Mostly scientifically accurate, but included some misconceptions or only partially explained antibiotic resistance. Student added some but not all of the information needed to explain the phenomena of antibiotic resistance. |
| 3 | Scientifically accurate and explained the concept of antibiotic resistance without any misconceptions. An ideal response mentioned how the microbes aren’t responding to the drugs using key terms or concepts such as - evolution, selection, mutation, and resistance. Explaining key terms without using them is okay. |

**Supplemental Table 5.** Rubric for scoring question 5: How would you explain antibiotic resistance to a fellow student in this class?

| Code | Description |
|---|---|
| Individual vs. Population (distinguish) | Made an explicit distinction between processes happening at the individual and group level (such as colonies, species, organisms, populations), OR stated that individuals can or can't do something, implying that it must happen at the group level. |
| Random | Mentioned that the mutation is developed by accident or the microbes don't have control over the development of the mutation. Stated that mutation is **not intentional** or a conscious decision. |
| Purposeful | Stated that the mutation is needed (or that yeast 'have to' acquire a mutation) to survive or to make a microbe(s) resistant. Implied that the mutation happens for a reason. Allowed responses that do not indicate it's random (e.g. 'individual microbes will develop resistance…'). |
| Natural selection | Made a reference to differential survival and reproduction. 'Pass on' or 'around' the group is allowed. Reference to survival needed to be related to differential survival vs. individual survival. Did not allow 'evolve' or 'adapt' without additional clarification of how these processes occur. |

**Supplemental Table 6.** List of code descriptions for question 6: [Explain why or why not] individual microbes develop mutations in order to become resistant to an antibiotic and survive.

| | Code | p-value |
|---|---|---|
| **Q1** | DNA | 0.99 |
| | **Code**** | **0.009** |
| | Make | **0.0726** |
| | Heredity | 0.6915 |
| | Trait | 0.2927 |
| **Q2** | Inherited | 0.6403 |
| | Phenotypic | 0.8329 |
| | Genotypic | 0.2736 |
| | Variation | 0.4448 |
| | Evolution | 0.1593 |
| **Q3** | **Adapt**** | **0.0019** |
| | Vague adapt | 0.181 |
| | Selection | 0.3039 |
| | Survive | 0.8798 |
| | Heredity | 0.0718 |
| | **Slow**** | **0.0121** |
| | **Mutation*** | **0.0253** |
| | Species | 0.5107 |
| | **Naive*** | **0.0313** |
| **Q4** | Change | 0.79 |
| | Trait | 0.2434 |
| | Heritable | 0.8884 |
| | Selection | 0.4858 |

| | | |
|---|---|---|
| | Essential for evolution | 0.6996 |
| | DNA | 0.6039 |
| | Adaptation | 0.9297 |
| | **Variation*** | **0.0399** |

**Supplemental Table 7.** Results of t-tests to test for significant differences between pre and post codes from questions 1, 2, 3, and 4.
