## Supplemental Text 2 for "yEvo: a modular eukaryotic genetics and evolution research experience for high school students"

### Azole Resistance Module 1
### Evolution of Drug Resistance

#### GOALS

1. Gain familiarity with sterile technique
2. Master skills of yeast culture by carrying out experiments for an extended period of time
3. Observe evolution in action by seeing the effects of selection pressure on yeast growth

#### OVERVIEW

**Evolution** is change in a population of organisms over time due to natural selection. Evolution acts on a population that has different characteristics, which affect their ability to survive and reproduce in specific conditions, called **selective pressures**. Different characteristics that can be selected upon are the result of mutations, which may exist in the starting population or arise over time. Evolution can be observed and measured using a laboratory technique called **experimental evolution**, in which organisms are grown under defined selective pressures. These can include growing bacteria in the presence of an antibiotic to study mutations that contribute to drug resistance, or growing yeast in acidic media to learn what mutations could improve fermentation of acidic foods. Mutations occur spontaneously, and some rare mutations can provide a fitness advantage. These rare mutations can improve growth under selective pressure, and thus will increase in frequency due to natural selection, sometimes to the point that all surviving organisms in the population contain these mutations.

The budding yeast ***S. cerevisiae*** is an ideal organism to use for experimental evolution because there are many resources available to study its genetics that help scientists interpret the results of experiments. *S. cerevisiae* is also used in many areas beyond the laboratory, where it naturally encounters selective pressures. Yeast used in bread making must be able to grow in high salt environments, and yeast used for industrial processes must be able to obtain energy from diverse fuel sources.

In these experimental evolutions, you will explore how yeast adapts to high concentrations of **clotrimazole**, the active ingredient in the anti-fungal FungiCure. Azole drugs like clotrimazole are one of the most commonly used antifungals in medicine and agriculture, and azole resistance is a growing global health crisis. Using experimental evolution, we can identify genetic factors that contribute to azole resistance. A better understanding of azole resistance can lead to new ways to treat azole-resistant fungal infections.

Every few days, you will transfer yeast in the presence or absence of FungiCure. Without FungiCure, the yeast should always grow well. With FungiCure, growth should be initially poor, but improve over time as mutant yeast arise in the cultures. After you have evolved FungiCure-tolerant yeast, you will compare their fitness with other groups' yeast to compare their ability to grow in FungiCure (Module 3-4). Some of your yeast will also be frozen and shipped to a university laboratory so they can be sequenced, and determine what mutations arose in your yeast that could contribute to their increased tolerance of caffeine (Module 2).

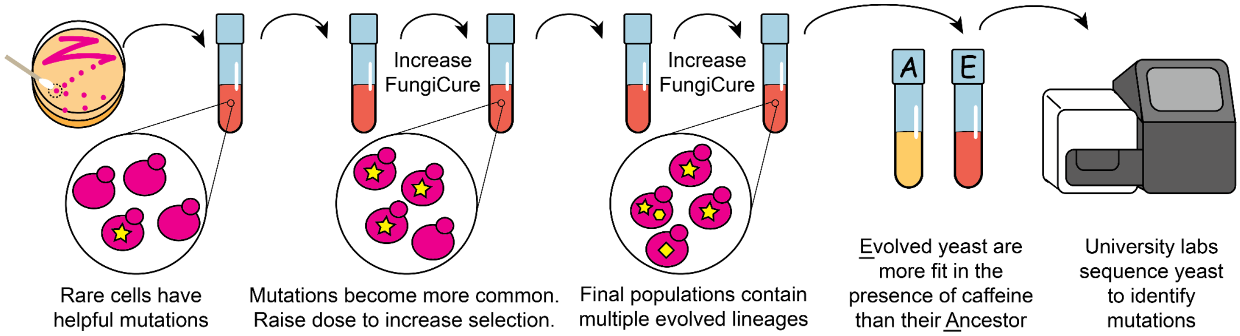

**Figure 1: Experimental evolution procedure overview.** *S. cerevisiae* with colored pigments are grown in the presence of FungiCure (clotrimazole). Random mutations arise, and beneficial mutations increase in frequency due to selective pressure as the dose of FungiCure is increased. The final population should be more fit than the ancestor in the presence of FungiCure; those yeast are sent to university labs for sequencing.

#### GLOSSARY

- Clotrimazole: An azole antifungal. Inhibits synthesis of ergosterol, a key membrane component and the fungal equivalent of cholesterol. Clotrimazole is the active ingredient in the FungiCure spray used in this experiment.
- Evolution: Change in a population of organisms over time due to natural selection. Evolution acts on a population that has different characteristics, which affect their ability to survive and reproduce under selective pressures. These different characteristics arise from mutations, which may exist in the starting population or arise over time.
- Fitness: A measure of an individual's reproductive success in a specific environment.
- G418: Geneticin; an antibiotic commonly used in laboratory experiments. Yeast utilized in this protocol are resistant to G418 due to a plasmid they carry, which also gives them their distinctive color thanks to additional genes on the plasmid that encode pigment production pathways. G418 is necessary for maintenance of the plasmid and additionally helps to prevent contamination.
- *Saccharomyces cerevisiae*: budding yeast, also known as baker's yeast, a unicellular fungus used in food production, industrial processes, and research.
- Selection pressure: An environmental condition that favors some genotypes in a population over others.
- YPD: A standard rich yeast medium named for its three ingredients: Yeast extract, Peptone, and Dextrose. Also referred to as YEPD.

#### TIME ESTIMATE

When growing yeast in an incubator, we recommend conducting 2-3 transfers per week and carry out the evolution for a minimum of 5 weeks. 5-10 weeks is ideal for the evolution, and allows time for additional modules that measure the fitness of evolved yeast to be carried out after the evolutions are stopped. When growing yeast at room temperature, we recommend conducting 1-2 transfers per week and carry out the evolution for 10-14 weeks.

#### MATERIALS AND EQUIPMENT

Yeast strains

- Ancestral *S. cerevisiae* strains carrying different pigment expression plasmids

Equipment

- Culture tubes (2 per group per transfer)
- P20 and P200 micropipettes and tips (for instructor to make FungiCure dilutions)
- Biohazard waste disposal bin, or a bucket of 10% bleach solution

Consumables

- YPD + G418 liquid media (4mL per group per transfer)
- FungiCure Intensive Maximum Strength Spray (1% clotrimazole in 70% isopropanol)
- Sterile swabs, sterile inoculating loops, or sterile inoculating sticks (2 per group per transfer)

For storage of yeast samples for sequencing (Module 2)

- Cryovials with 0.5mL 50% sterile glycerol (1 per group for every 1-3 weeks)
- Freezer, preferably NOT frost-free (so it does not go through temperature cycles)
- P1000 micropipette or other pipettor to measure 0.5mL
- Fine-tipped permanent marker

*Optional*

- *30ºC incubator*
- *Test tube roller drum or shaking platform*
- *70% isopropanol (for diluting FungiCure to add directly to students' tubes)*

#### SAFETY

1. The colored yeast strains are genetically modified and thus considered biohazardous waste. Dispose of inoculating sticks in a biohazard bin, or decontaminate by placing in 10% bleach for 20 minutes before throwing away. Liquid waste containing yeast should

be collected and decontaminated using bleach at a final concentration of 10% for 20 minutes before pouring down a drain.

2. **SPILLS:** If there are any spills of the yeast cultures, they should be blotted with paper towels by placing the towels over the spill. Then the towels should be sprayed with 10% bleach and left for 10 minutes before cleaning up.
3. **CONTACT EXPOSURE:** If you spill yeasts on your hands, wash then thoroughly with hot soapy water. If the yeast splash into your eyes, flush them with warm running water. Yeast splashed on clothing should be blotted and washed with soapy water.
4. **CONTAMINATION:** If you notice that the color of your yeast culture has changed or that the culture has gone moldy – **DO NOT OPEN the tube**. The tube has become contaminated and the entire tube (including the liquid) should be immersed in 10% bleach. Tubes should be left for 20 minutes before disposal of the liquid down the sink and tubes into the trash or proper glass disposal.
5. **FUNGICURE:** FungiCure Intensive Maximum Strength Spray (1% clotrimazole) is available commercially over the counter. The small quantities used in our experiments **do not** pose a significant health hazard. The FungiCure is dissolved in 70% isopropanol which is extremely **FLAMMABLE**: keep away from flames and other sources of ignition. Isopropanol may cause skin irritation and severe eye **irritation: wear protective eyewear when pipetting FungiCure**. If you spill FungiCure on your hands, wash them thoroughly with warm soapy water. If FungiCure splashes into your eyes, flush them with warm running water.

#### BEFORE THE LAB: Week 1

1. Plan out how the timing of activities will fit with your class schedule. Yeast grow most robustly at 30ºC. They can be grown at room temperature as well but will grow more slowly. When growing at 30ºC, transfers can occur 2-3 times per week. When growing at room temperature, transfers can occur 1-2 times per week.
2. Streak ancestral strains with different colored plasmids onto YPD + G418 agar media 3-5 days before the intended start of the lab and place in 30ºC incubator to grow. If you have a plate with the strains saved, that is sufficient and you can proceed to step 3.
3. Inoculate tubes with different colored strains 1-2 days before the lab. You can prepare one per strain, or one tube for each group. Place 2mL of YPD + G418 media into a tube, use a sterile swab to pick up a colony of yeast, swirl in the media, and put at 30ºC to grow.
4. Prepare tubes with media. Each group will need 1 tube with 2mL of YPD + G418 (labeled "0µM") and 1 tube with 2mL of YPD + G418 + 10µM clotrimazole (labeled "2.5µM").
   a. To make 2.5µM clotrimazole media, add 3.88µL FungiCure (1% clotrimazole) to 50mL YPD + G418 (1:12,800 dilution).

#### PROTOCOL: Week 1

1. Take two two tubes filled with YPD + G418 growth medium, one without FungiCure and one with 2.5µM.

    *The negative control is **crucial** for the students to observe the evolution of resistance. Once the tube with FungiCure appears to be growing as well as the control (and by comparing to their previously recorded pictures or notes), the students should double the concentration of FungiCure.*

2. Label both tubes with your group name and date, and the concentration of FungiCure ("0µM" or "none", and "2.5µM")
3. Using a sterile swab, dip it into the yeast culture that you have been provided with.
4. Transfer the damp swab (the liquid in the cotton bud will have millions of yeasts stuck to it) to the tube for no FungiCure.
5. Mix the growth media in the new tube with the swab – remove the swab and dispose of with biohazardous waste (in a biohazard bin or a pot of 10% bleach).
6. Using another sterile swab, dip it into the yeast culture that you have been provided with.
7. Transfer the damp swab to the tube with 2.5µM FungiCure.
8. Mix the growth media in the new tube with the swab – remove the swab and dispose of with biohazardous waste.
9. Record observations about your cultures as instructed in the "Questions: Week 1" section below.
10. Place your tubes in a rack to go into the 30$^{o}$C incubator.

#### QUESTIONS: Week 1

1. As a class, come up with a hypothesis about how the yeast will grow over the course of the evolution experiment. Record that hypothesis.
2. Look at the yeast cultures. You can take a picture and describe what the yeast culture looks like. Is it see-through or opaque (cloudy/muddy)? Compare the tubes that you just inoculated to the tubes that you were given. Write down what you observe.
3. Record your group name and the color of your yeast strain at the top of Table 1. This is important since you will use your name to label your tubes and the yeast that are frozen for sequencing. Make sure that this name is unique and not used by any other groups! (Your instructor may assign you a group name.)

#### BEFORE THE LAB: Subsequent weeks

1. Prepare tubes with media. Each group will need 1 tube with 2mL of YPD + G418 (labeled "0µM") and 1 tube with 2mL of YPD + G418 + FungiCure. The concentration of FungiCure will depend on the growth students observe. You can prepare tubes with different concentrations of caffeine, or make dilutions of FungiCure in 70% isopropanol that can be directly added to students' tubes.
    a. For making different concentrations in media, add the µL of FungiCure indicated in the table below to 50mL YPD + G418 media. Scale as needed.
    b. For making different stock solutions (100x concentration), add the µL of FungiCure indicated in the table below to 70% isopropanol to reach a final volume of 500µL. Then add 20µL of 100x FungiCure stock to 2mL YPD + G418 in students' tubes.

| **Final µM** | **2.25** | **4.5** | **9** | **18** | **36** | **72** | **144** |
|---|---|---|---|---|---|---|---|
| µL FungiCure | 3.88 | 7.76 | 15.5 | 31.0 | 62.1 | 124 | 248 |

#### PROTOCOL: Subsequent weeks

1. Retrieve your yeast cultures.
2. Check for contamination: The yeast should have settled to the bottom of your tube, and the media above it remain mostly clear. If the media is cloudy, it could indicate bacterial contamination. Inform your instructor and consider re-starting your culture from a previous week's tube (or from the original stock if this is the first transfer).
3. Swirl your tubes around so that the yeast are suspended in the liquid.
4. Record observations as instructed in Table 1.
5. Optional: take a picture of your yeast. Look at the photo from last week – notice any changes? Use this information to help you complete Table 1.
6. Decide if you will increase the concentration of FungiCure. If the growth of yeast with FungiCure looks similar to growth of yeast without FungiCure, double the concentration of FungiCure you will use. Record this in Table 1.
7. Transfer the "no FungiCure" culture
    a. Obtain a tube with YPD + G418 media. Label it with group name, date and "0µM".
    b. Using a sterile swab, dip it into the yeast culture labelled "0µM" that you grew.
    c. Transfer the damp swab into the new tube that you have prepared in step a.
    d. Mix the growth media in the new tube with the swab – remove the swab and dispose of in biohazardous waste.

8. Transfer the "FungiCure" culture.
    a. Obtain a tube with YPD + G418 media. If you decided to increase the concentration, choose the next highest dose of FungiCure, doubling the concentration you used last week. Label it with group name, date and the concentration of FungiCure.
    b. Using a sterile swab, dip it into the yeast culture with FungiCure that you grew.
    c. Transfer the damp swab into the new tube that you have prepared in step a.
    d. Mix the growth media in the new tube with the swab – remove the swab and dispose of in biohazardous waste.
9. Place your new tubes in a rack to go into the 30ºC incubator.
10. If it is the end of the week, proceed to "Storing yeast for sequencing"
11. Place your old tubes (that you transferred from) into a rack that stays at room temperature. This is in case anything goes wrong with your new cultures, you can return to these tubes instead of starting over from the beginning!

#### PROTOCOL: Storing yeast for sequencing

*This can be completed by each group, or by the instructor / with the instructor's assistance.*

1. At the end of the week, obtain a cryovial containing 0.5mL 50% glycerol
2. Label the side of the vial with your group name, yeast color, and the date
3. Add 0.5mL of your yeast culture that has been evolved in FungiCure (not the 0µM FungiCure control strain)
4. Invert the tube 5 times to mix together
5. Place the tube in the freezer or collection area as instructed

*These stocks can also be used if a group encounters contamination or loses their sample, and does not have one from a previous week. Use a toothpick or pipette tip to scrape out a bit of the glycerol stock, patch it onto a YPD + G418 agarose plate, spread out, and place in 30ºC incubator for 2-3 days. Pick up multiple colonies with a swab to re-start the culture, to recapture some of the heterogeneity of the frozen population.*

#### TABLE 1: Observations and notes - example

Use this table to record how your yeast is growing in the presence of FungiCure, when you change the concentration, and anything else that occurs during the experiment you need to record.

Example: You have been growing your yeast in 4.5µM FungiCure, and it was of comparable density to the yeast in no FungiCure on 4/3. You decide to transfer it into 9µM FungiCure. When you check your tubes on 4/7, there is no growth in the 9µM FungiCure! You conclude that the yeast was not yet fit enough to survive in 9µM, so you go back to your tube from 4/3 and transfer some to 4.5µM FungiCure to continue to grow. Your notes should look something like this:

| Date | FungiCure concentration | Observations and notes | New FungiCure concentration |
|---|---|---|---|
| 3/31 | 4.5µM | Culture is very cloudy but not as much as no FungiCure tube. | 4.5µM |
| 4/3 | 4.5µM | Growing like no FungiCure culture! Ready to raise concentration. | 9µM |
| 4/7 | 9µM | No growth. Went back to tube from 4/3 to re-start. | 4.5µM |

**TABLE 1: Observations and notes**

**Group name: ____________________________________ Yeast color: ________________**

| Date | FungiCure concentration | Observations and notes | New FungiCure concentration |
|---|---|---|---|

| Date | FungiCure concentration | Observations and notes | New FungiCure concentration |
|---|---|---|---|

| Date | FungiCure concentration | Observations and notes | New FungiCure concentration |
|---|---|---|---|

#### QUESTIONS: Final day

1. What was the highest concentration of FungiCure at which you were able to grow your yeast?
2. Do your results support the hypothesis that you developed as a class in the first week of the experiment? Explain how your data does or does not support the hypothesis.
3. Compare your final concentration and the density (cloudiness) of your evolved yeast culture to other groups. Do you think that your yeast would be more or less capable of growing in a high concentration of FungiCure than other groups' yeast? Support your answer with observations of the yeast cultures.
4. Where did you encounter challenges in the evolution experiment? Did you ever have to go back to a previous culture? How do you think this affected the fitness of your final evolved yeast in FungiCure? Comparing your experiences and yeast cultures with other groups' may help you answer this question.
5. If you had time to do another evolution experiment, what condition would you want to expose yeast to?
