## Supplemental Text 3 for "yEvo: a modular eukaryotic genetics and evolution research experience for high school students"

### Azole Resistance Module 2
### Genome Sequence Analysis

#### GOALS

1. Learn how to use online genetics analysis tools.
2. Identify different types of mutations and their effects.
3. Form specific hypotheses on how mutations may contribute to altered fitness.

#### OVERVIEW

The evolution experiments in Module 1 select for yeast mutants that are more resistant to clotrimazole (the active ingredient in FungiCure). These mutants will possess a few specific changes to their genetic code that cause them to be resistant to clotrimazole. After Module 1, we isolated yeast from each experiment and sequenced their genomes to identify their mutations.

In this module, you will receive the sequence of one of the genes found to be mutated in your evolved yeast. Using online tools, you will identify the gene and type of mutation, and research its function in yeast. From this information, you will form hypotheses about how this mutation may contribute to the azole resistance of your evolve strain.

#### MATERIALS AND EQUIPMENT

- Computer and internet access for each participant
- .xlsx mutation file provided by university lab partner, which contains a list of mutations from a strain evolved in Module 1

#### INTRODUCTION

Your yeast have had the genetic code of their entire genome determined! The yeast genome is 12 million letters long. Your yeast likely had between 1 and a handful of single changes to its genome sequence. We have sent you a list of mutations that your yeast possessed, as well as an altered sequence from your yeast. We will walk through how to build hypotheses about what these mutated genes are doing.

Yeast have thousands of genes that each are a blueprint for making a bit of cellular machinery called a protein. These machines come together in intricate ways to form assembly lines (like the ergosterol "pathway"), rigid support structures (like actin), export systems (like the pump *PDR5*), or master regulators of specific processes (like the transcription factor *UPC2*). The goal of this activity is to think through how the DNA changes in your evolved yeast will impact your genes' protein and ultimately make your yeast more resistant to the FungiCure. Look at **Figure 1** for some examples of genes and pathways that are known to be affected by FungiCure (clotrimazole is the active ingredient in FungiCure).

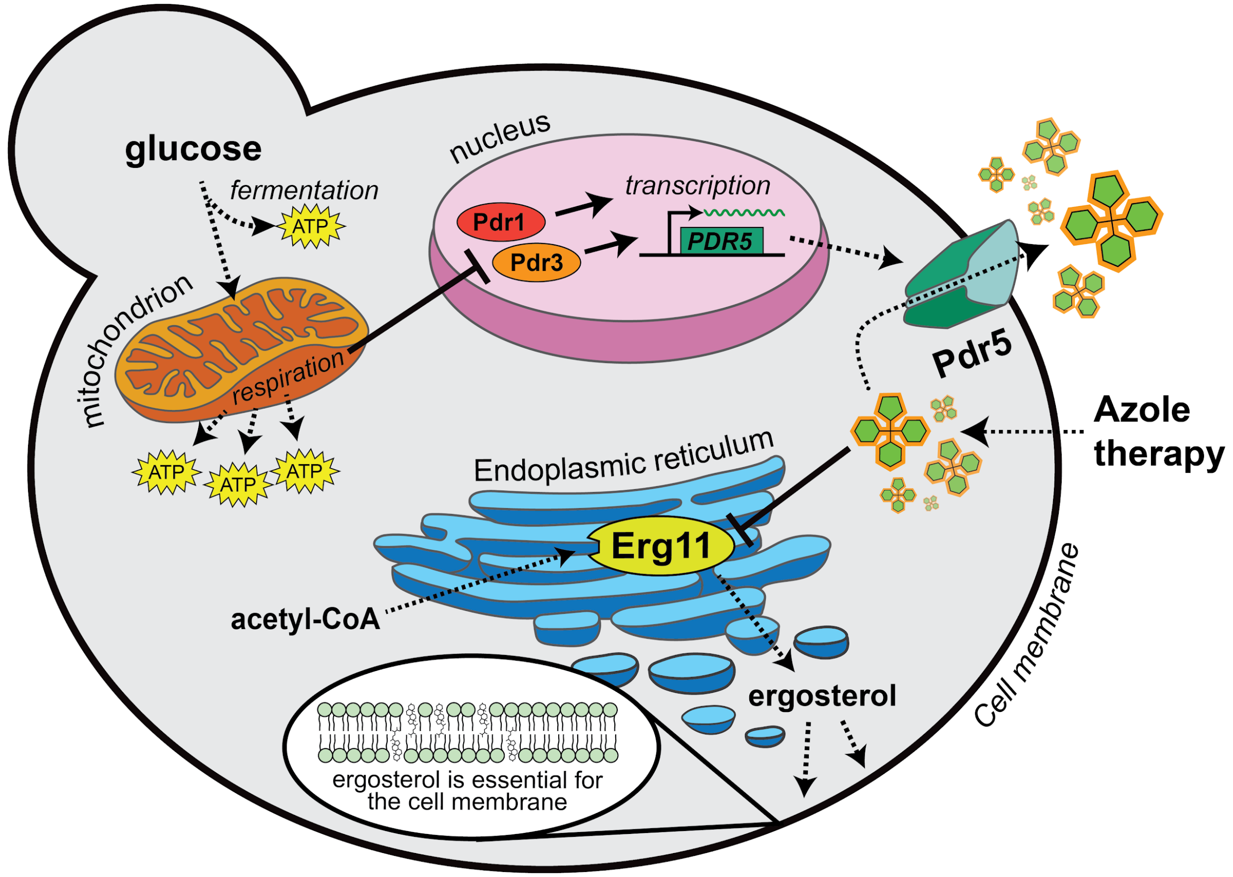


**Figure 1:** Pathways affected by azole therapy clotrimazole, the active ingredient in FungiCure. Clotrimazole inhibits the activity of Erg11 to reduce the production of ergosterol, an essential membrane sterol. Proteins and processes that are important for clotrimazole resistance are illustrated. Mutations in genes *PDR1* and *PDR3* or loss of mitochondrial respiration result in an increase in the expression of *PDR5* and the active removal of clotrimazole from the cell. Duplications of *ERG11* compensate for inhibition by clotrimazole. Other pathways not featured here can allow yeast to survive with less ergosterol.

Now we will use this information to learn more about how your yeast evolved! You have a file named mystery_gene#.fas (with some number in place of the #) that contains the sequence of a gene from your yeast that we think might have been important for this resistance. We will work through the steps to determine what this mystery gene is and its function. We will begin by looking at the DNA sequence of your evolved yeast to see how it has changed, and how this will change the gene's protein sequence. We will then use a database to figure out what your gene does. This information can be used to understand how your yeast evolved to become resistant to FungiCure.

For a refresher on how a gene's DNA sequence is used to produce a protein, we recommend checking out Khan Academy's Central Dogma section, particularly the article on the genetic code.

#### TASK 1: Identify your mutation

Before we begin our analysis, let's look at your mutation file.

Your mutation is one of 4 types of genetic changes:

1. **Synonymous mutations** are DNA changes that do not change an amino acid. You will not be able to see these in the protein alignment. They are sometimes referred to as silent mutations because they do not impact a gene's protein.
2. **Missense mutations** change an amino acid in a gene's protein. They may change the function of that protein.
3. A **nonsense mutation** results in an early stop in the gene. You will notice the * in the sequence of your query if you have a nonsense mutation. None of the amino acids that come after this stop will be added to the protein, so the resulting protein will be shorter than the original subject (sbjct) version.
4. A **frameshift mutation** is when one or a few nucleotides are inserted or deleted and results in a change in the reading frame of a protein, shifting how the code of the gene is read. Nearly all the amino acids after the mutation will be changed, and this frequently leads to early stops, like in the example blastx alignment above.

#### TASK 1 QUESTIONS

1. What change(s) can you see in your evolved yeast's DNA?
2. Record how your mutation impacted your amino acid sequence. What was the original codon, and what is the codon after the mutation (see Resource 1). What was the original amino acid, and what is the amino acid after the mutation occurred?
3. Which of the 4 types of mutations (above) matches the one you see? Do you think it will severely affect the function of the protein? Why or why not?

#### TASK 2: Learn about the gene affected by your mutation

We will now move onto the *Saccharomyces* Genome Database (SGD) website to learn more about our genes. SGD contains information from thousands of published studies on yeast, which has been curated to make it easier to find information about a specific gene.

https://www.yeastgenome.org/

For this example, I'm going to show you the page for a gene called *ERG11*. This gene encodes instructions for making an enzyme that plays a role in making ergosterol. Ergosterol is like cholesterol, and it helps cell membranes to stay structurally sound. Clotrimazole, the drug you worked with in Module 1, stops this enzyme from working, which ultimately harms the cell's membrane and prevents it from growing.

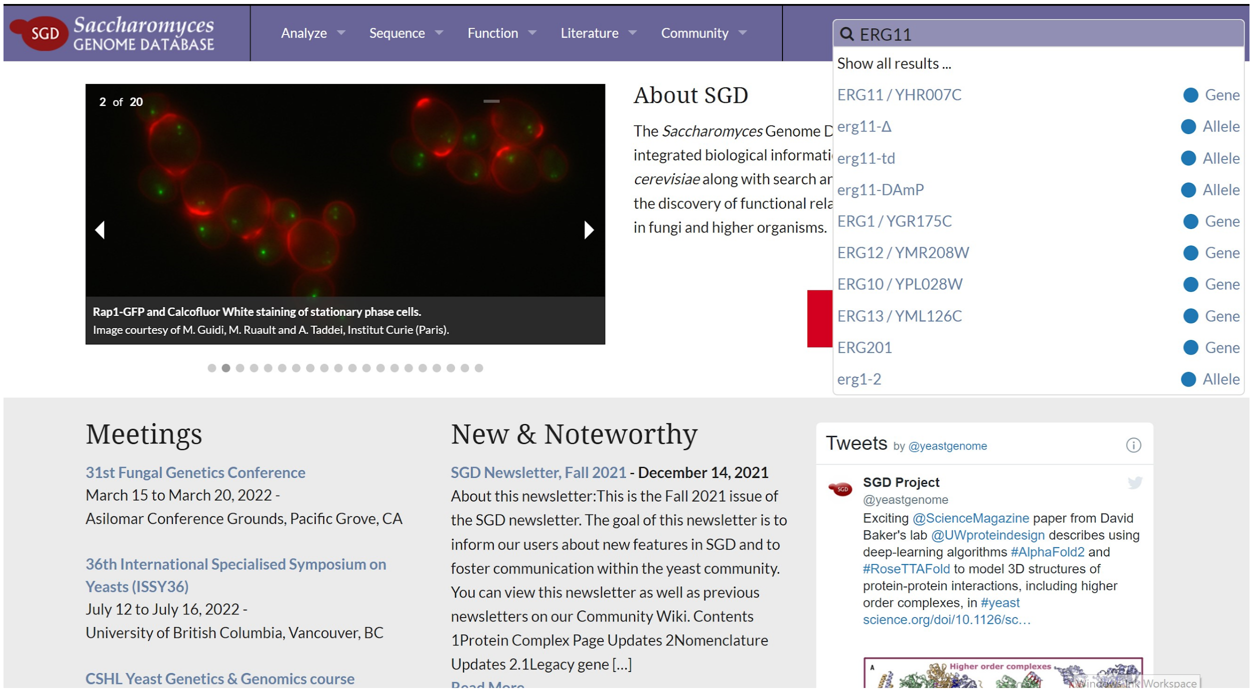


First let's go to the *ERG11* page. You can do this by doing a google search for "ERG11 SGD", or by entering "ERG11" in the search bar at the top right of the SGD homepage as shown above.

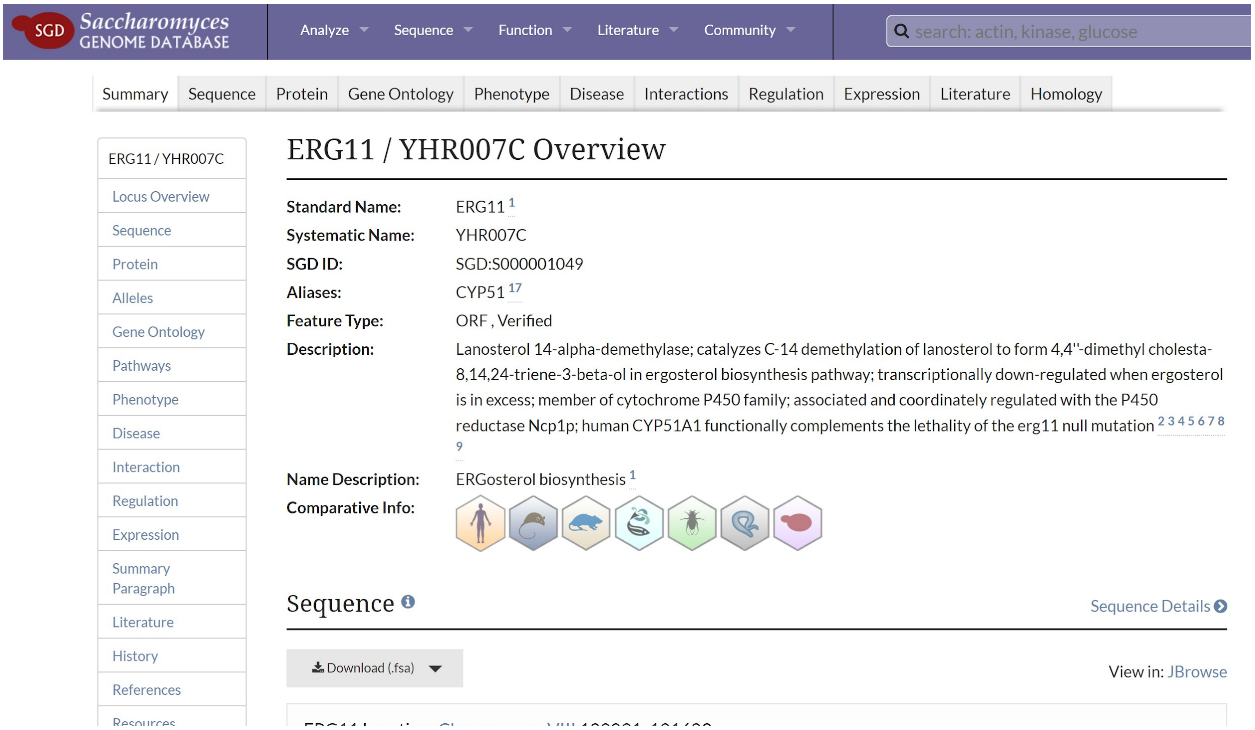

This page contains a ton of info organized into sections that are listed in light blue along the left side of the page. The sections I like to use are *Locus Overview*, *Summary Paragraph*, and *References*. The others are great, but for now you can ignore them.

When you first enter the page you should see the Locus Overview, displayed as "ERG11 / YHR007C Overview". The Description section will give you a short description of what is known about the gene. It's extremely dense, and you will encounter a lot of unfamiliar terminology. The trick is to look through for terms that you recognize that are related to the clotrimazole drug from Module 1.

In this case, the term "ergosterol" is a great clue. In Module 1 we worked with a drug that inhibits ergosterol synthesis, and here we have a gene that is important for synthesizing ergosterol. The chances of a gene with this role being randomly mutated in your strain is very low, so there's a good chance this mutation was selected during the evolution experiment.

The Comparative Info section can be cool to click through. It shows icons that represent other organisms that possess this gene. Most of the genes that make ergosterol in yeast also make cholesterol in mammals, so this section includes an icon for humans, mice, and rats. If you click on the icon you'll be taken to a different database that has information about how this gene works in those organisms. You don't need to do that now, but it's worth checking out if you finish early.

Now navigate down to Summary Paragraph.

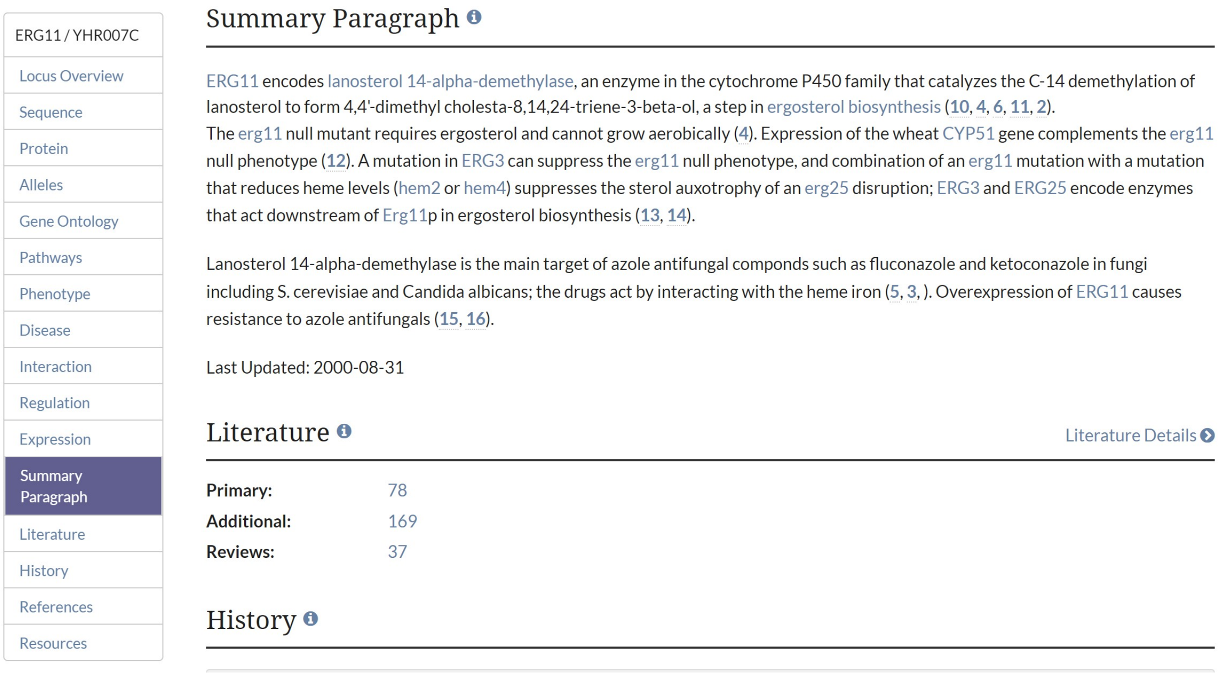
ERG11 / YHR007C
Locus Overview
Sequence
Protein
Alleles
Gene Ontology
Pathways
Phenotype
Disease
Interaction
Regulation
Expression
Summary Paragraph
Literature
History
References
Resources

Summary Paragraph

ERG11 encodes lanosterol 14-alpha-demethylase, an enzyme in the cytochrome P450 family that catalyzes the C-14 demethylation of lanosterol to form 4,4'-dimethyl cholesta-8,14,24-triene-3-beta-ol, a step in ergosterol biosynthesis (10, 4, 6, 11, 2).
The erg11 null mutant requires ergosterol and cannot grow aerobically (4). Expression of the wheat CYP51 gene complements the erg11 null phenotype (12). A mutation in ERG3 can suppress the erg11 null phenotype, and combination of an erg11 mutation with a mutation that reduces heme levels (hem2 or hem4) suppresses the sterol auxotrophy of an erg25 disruption; ERG3 and ERG25 encode enzymes that act downstream of Erg11p in ergosterol biosynthesis (13, 14).

Lanosterol 14-alpha-demethylase is the main target of azole antifungal componds such as fluconazole and ketoconazole in fungi including S. cerevisiae and Candida albicans; the drugs act by interacting with the heme iron (5, 3, ). Overexpression of ERG11 causes resistance to azole antifungals (15, 16).

Last Updated: 2000-08-31

Literature
Literature Details

| | |
|---|---|
| Primary: | 78 |
| Additional: | 169 |
| Reviews: | 37 |

History

The summary paragraph is a longer description of what we know about this gene. It's usually in more of a narrative format and can be easier to read (but not always).

Finally, check out References.

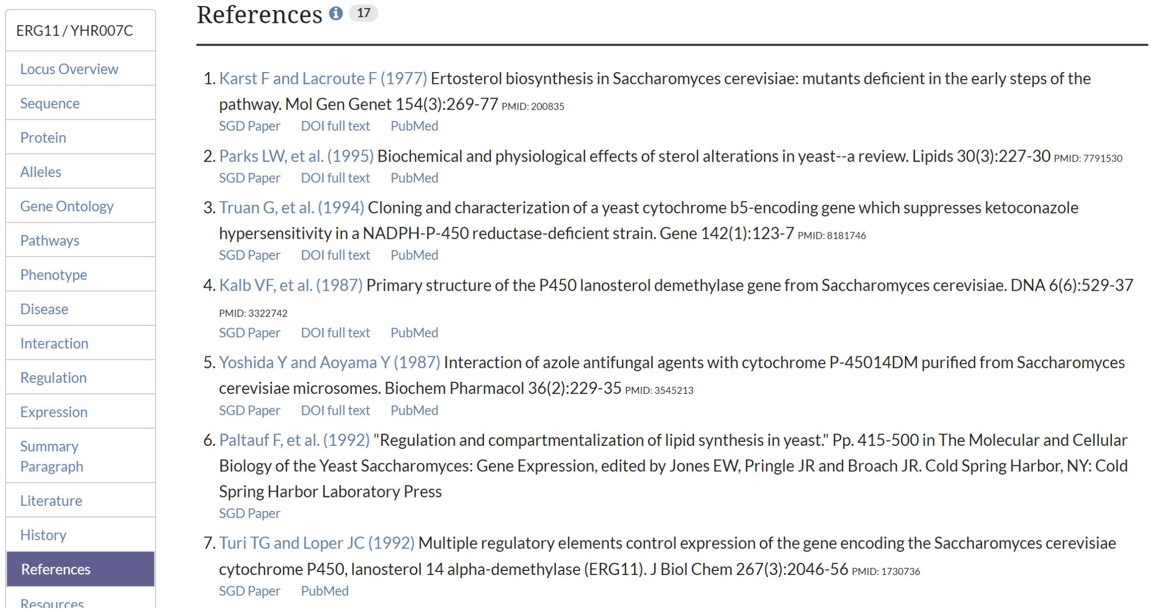


This section lists scientific publications that include information about the gene. The SGD page for each gene is made by combing through publications like these to find useful information about what the gene does.

You may have noticed light blue numbers in the sections above. These numbers are citations for information and correspond to publication in this list. If you saw information above that seemed useful, you can click on the “SGD Paper” link under a publication to find a freely-accessible text of the article.

Skimming publication titles in the references section may lead you to more information about what your gene does.

Now, go to the SGD page for one of the genes in your list and answer the questions below.

#### TASK 2 QUESTIONS

1. Find one fact about your gene and record it.
2. What is the function of your gene?
3. Did you notice any keywords related to clotrimazole for your gene?
4. Can you think of a way your gene’s function could be related to azole drug resistance?

RESOURCE 1

**Codon table**

Second Letter

| 1st letter | U | | C | | A | | G | | 3rd letter |
|---|---|---|---|---|---|---|---|---|---|
| U | UUU | Phe | UCU | | UAU | Tyr | UGU | Cys | U |
| | UUC | | UCC | Ser | UAC | | UGC | | C |
| | UUA | Leu | UCA | | UAA | Stop | UGA | Stop | A |
| | UUG | | UCG | | UAG | Stop | UGG | Trp | G |
| C | CUU | | CCU | | CAU | His | CGU | | U |
| | CUC | Leu | CCC | Pro | CAC | | CGC | Arg | C |
| | CUA | | CCA | | CAA | Gln | CGA | | A |
| | CUG | | CCG | | CAG | | CGG | | G |
| A | AUU | | ACU | | AAU | Asn | AGU | Ser | U |
| | AUC | Ile | ACC | Thr | AAC | | AGC | | C |
| | AUA | | ACA | | AAA | Lys | AGA | Arg | A |
| | AUG | Met | ACG | | AAG | | AGG | | G |
| G | GUU | | GCU | | GAU | Asp | GGU | | U |
| | GUC | Val | GCC | Ala | GAC | | GGC | Gly | C |
| | GUA | | GCA | | GAA | Glu | GGA | | A |
| | GUG | | GCG | | GAG | | GGG | | G |

**RESOURCE 2**

**Amino acid codes table**

| Amino Acid | 3-Letter Code | 1-Letter Code |
| --- | --- | --- |
| Alanine | Ala | A |
| Cysteine | Cys | C |
| Aspartic acid or aspartate | Asp | D |
| Glutamic acid or glutamate | Glu | E |
| Phenylalanine | Phe | F |
| Glycine | Gly | G |
| Histidine | His | H |
| Isoleucine | Ile | I |
| Lysine | Lys | K |
| Leucine | Leu | L |
| Methionine | Met | M |
| Asparagine | Asn | N |
| Proline | Pro | P |
| Glutamine | Gln | Q |
| Arginine | Arg | R |
| Serine | Ser | S |
| Threonine | Thr | T |
| Valine | Val | V |
| Tryptophan | Trp | W |
| Tyrosine | Tyr | Y |
