## Supplemental Text 4 for "yEvo: a modular eukaryotic genetics and evolution research experience for high school students"

### Azole Resistance Module 3
### Fitness – Growth Inhibition

#### GOALS

1. Demonstrate that strains evolved in the presence of an antifungal are more fit than their ancestors in that environment
2. Demonstrate that independent evolved strains may differ in their fitness in the evolved environment

#### OVERVIEW

The goal of this lab is to provide visual evidence of evolution. You will use a Minimum Inhibitory Concentration (MIC) assay to quantitatively determine the resistance of evolved yeast from Module 1.

In an MIC assay, a microbe is exposed to several doses of a drug to determine the minimum dose that completely inhibits growth. In this instance, we will expose yeast from the Module 1 evolution experiments to several doses of FungiCure. The MIC is the lowest dose of a compound - here, FungiCure - that completely inhibits growth of your yeast strain (**Figure 1**). Yeast that evolved in the presence of FungiCure should be able to withstand higher doses than their ancestors, and therefore have a higher MIC. Not all experiments reach the same fitness, so results may vary!

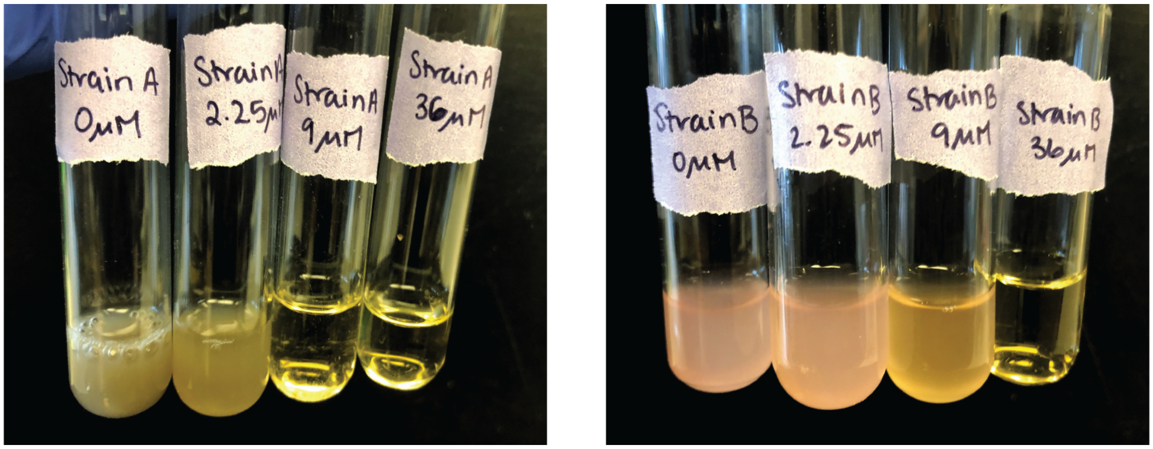


**Figure 1: Minimum Inhibitory Concentration (MIC).** Two yeast strains (A, with a gray pigment, and B, with a pink pigment) were evolved in the presence of FungiCure. After the end of the evolution experiment, a colony from each strain was grown overnight and then inoculated into YPD + G418 media containing different doses of FungiCure (clotrimazole). The lowest doses that completely inhibited growth for A and B were 9μM and 36μM, respectively. Therefore, the MIC of clotrimazole for Strain A is 9μM, and for Strain B is 36μM.

#### GLOSSARY

- Clotrimazole: An azole antifungal. Inhibits synthesis of ergosterol, a key membrane component and the fungal equivalent of cholesterol. Clotrimazole is the active ingredient in the FungiCure spray used in this experiment.
- Fitness: A measure of an individual's reproductive success in a specific environment.
- G418: Geneticin; an antibiotic commonly used in laboratory experiments. Yeast utilized in this protocol are resistant to G418 due to a plasmid they carry, which also gives them their distinctive color thanks to additional genes on the plasmid that encode pigment production pathways. G418 is necessary for maintenance of the plasmid and additionally helps to prevent contamination.
- MIC: Minimum Inhibitory Concentration. The lowest dose of a drug or other stressor that completely inhibits the growth of a microbe.
- Selection pressure: An environmental condition that favors some genotypes in a population over others.
- YPD: A standard rich yeast medium named for its three ingredients: Yeast extract, Peptone, and Dextrose. Also referred to as YEPD.

#### MATERIALS AND EQUIPMENT

Yeast strains

- Evolved and ancestral *S. cerevisiae* strains (from Module 1)

Equipment

- Pipettes: volume needs will vary based on implementation. You will likely need a P2-20ul, a P20-200ul, and a P200-1000ul or equivalent, as well as a 5ml serological pipette.
- Culture tubes
- Glass beads or plate spreader

Consumables

- YPD + G418 liquid media (at least 30ml per yeast strain per student)
- Sterile swabs, sterile inoculating loops, or sterile inoculating sticks
- YPD + G418 agar plate (to streak out starting strains)

Chemicals

- FungiCure spray with active ingredient clotrimazole

*Optional*

- *30°C incubator*
- *Test tube roller drum or shaking platform*
- *Vortex machine*

#### BEFORE THE LAB

1. Plan out how the timing of activities will fit with your class schedule. Yeast grow most robustly at 30ºC. They can be grown at room temperature as well but will grow more slowly. *We've included estimates for the time it'll take for your students' yeast to grow where applicable in italics*.
2. Make FungiCure media. Each competition performed will require 5ml each of a low and a high dose of FungiCure media. In our hands, the concentrations below work well, but we encourage you to experiment with additional doses as time and resources permit!
   - Low dose: 1:12,800x, 2.25µM (3.88ul of fungicure in 50ml of YPD + G418) inhibits growth the ancestral strains and was the starting concentration for the evolutions.
   - High dose: 1:3200x, 9µM (15.5ul of fungicure in 50ml of YPD + G418) prevents growth of the ancestral strains but not the evolved strains.
   - Very high dose: 1:800x, 36µM (62.1ul in 50ml of YPD + G418) prevents growth of the ancestral strains and several evolved strains, but some evolved strains will grow.
3. Streak evolved and ancestral strains onto YPD + G418 agar media at least 2 days before the intended start of the lab.

#### PROTOCOL

**Day 1:** Inoculate evolved and ancestral strains of yeast into separate tubes of medium.

1. Fill two test tubes with 5ml each of liquid YPD + G418. Label one "evolved" and the other "ancestor".
2. Use a sterile swab, inoculating loop, or inoculating stick to pick a colony of either evolved or ancestral yeast and inoculate it into its respective test tube.
3. Allow the yeast in these tubes to grow until you can no longer see through the liquid media. *When growing at 30ºC in a roller drum or shaking platform, this will take 1-2 days. When growing on a bench top at room temperature without shaking or rolling it will take 2-3 days. Yeast can be left longer than these amounts of time (up to a week) without worry*.

**Day 2:** Mix evolved and ancestral strains in media with or without FungiCure.

1. Fill one test tube each with 5ml of the following three media: YPD + G418; YPD + G418 + low dose of FungiCure; YPD + G418 + high dose of FungiCure; YPD + G418 + very high dose of FungiCure. Label these tubes with the media type used.
2. Examine the cultures you inoculated on Day 1. If yeast have settled at the bottom of the tube (pelleted), gently shake the tube until they are completely resuspended. The liquid sample of yeast (culture) should be dense enough that you cannot see through it, and the two cultures should be comparable in density.

3. **For each strain to be assayed:** Add 5ul of culture to each of the four test tubes you inoculated. *Incubate these as on Day 1.*

**Day 3:** Observe growth of yeast strains in various doses of FungiCure.

1. Examine the cultures you inoculated on Day 2. If yeast have settled at the bottom of the tube (pelleted), gently shake the tube until they are completely resuspended. The liquid sample of yeast (culture) should be dense enough that you cannot see through it, though the culture grown in the highest dose of FungiCure may appear less dense than the others.
2. Record the lowest dose that completely inhibits growth of each of your yeast strains. This is the MIC for that strain.

#### EXTENSION QUESTIONS

1. How do your results compare to those of other groups?
2. Do all yeast adapted to the same environment (evolved to survive in the same dose of FungiCure) have the same fitness in that environment?
