## Supplemental Text 5 for "yEvo: a modular eukaryotic genetics and evolution research experience for high school students"

### Azole Resistance Module 4
### Fitness – Competition

#### GOALS

1. Demonstrate that strains evolved in the presence of an antifungal are more fit than their ancestors in that environment
2. Demonstrate that independent evolved strains may differ in their fitness in the evolved environment

#### OVERVIEW

We have been carrying out experimental evolutions with lab strains of *S. cerevisiae* that express vibrant pigments and thus each have a distinct color. Because of these colors, relative abundance of each strain in a mixed culture can be determined by counting colony forming units (CFUs) and calculating the ratio of colors (**Figure 1**). This approach can be used to determine whether yeast with different colors are better adapted to a particular environment.

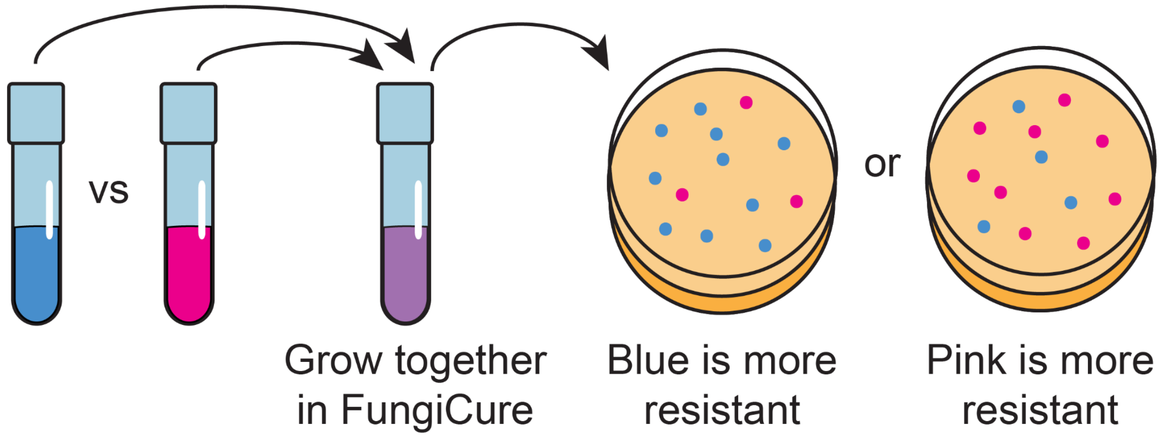


**Figure 1: Competition experiment overview.** Ratio of CFU colors on agar media indicates the relative abundance of each strain in a mixed culture.

We will use these colors in a competition experiment, which will allow us to determine which yeast from your experiments are best-adapted to the antifungal drug used in Module 1. To do this, you will first grow yeast from different timepoints in the evolution experiment and with different colors in a medium that does not contain the antifungal. You will then mix these strains in media containing varying concentrations of the antifungal so that they will compete for resources, You will then plate these mixed cultures onto agar media. After a few days of growth, you can count the ratio of colors on each plate to determine which strain “won” the competition in each concentration (**Figure 2**).

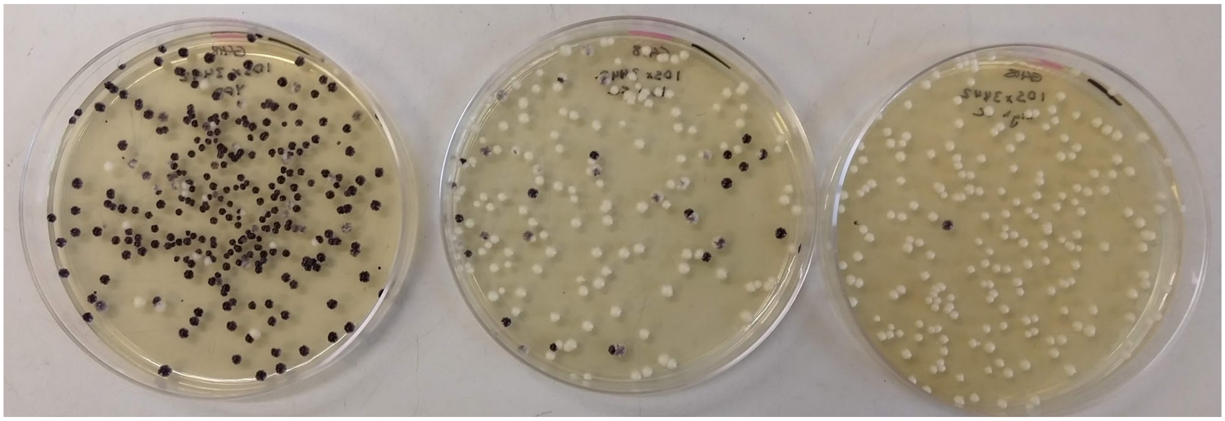

**Figure 2: Example of competition experiment outcome.** A black ancestral strain and a cream evolved strain were co-cultured in YPD + G418 media containing no FungiCure, a low dose, and a high dose for 24 hours. A 1:10,000 dilution of each culture was plated onto YPD + G418 agar media. Left plate: no FungiCure; middle plate: low dose (2.25µM); right plate: high dose (9µM).

#### GLOSSARY

- CFU: Colony Forming Unit; a cell that is capable of growing into a colony of cells when transferred from liquid to solid medium. CFUs are commonly used as a proxy for the number of viable cells in a liquid culture.
- Clotrimazole: An azole antifungal. Inhibits synthesis of ergosterol, a key membrane component and the fungal equivalent of cholesterol. Clotrimazole is the active ingredient in the FungiCure spray used in this experiment.
- Fitness: A measure of an individual's reproductive success. Note that in this experiment you will not be measuring fitness directly but will instead observe the outcome of competition between individuals with differing fitness in a given environment. Success in this competition is determined by the relative fitness between these individuals.
- G418: Geneticin; an antibiotic commonly used in laboratory experiments. Yeast utilized in this protocol are resistant to G418 due to a plasmid they carry, which also gives them their distinctive color thanks to additional genes on the plasmid that encode pigment production pathways. G418 is necessary for maintenance of the plasmid and additionally helps to prevent contamination.
- Selection pressure: An environmental condition that favors some genotypes in a population over others.
- YPD: A standard rich yeast medium named for its three ingredients: Yeast extract, Peptone, and Dextrose. Also referred to as YEPD.

#### MATERIALS AND EQUIPMENT

Yeast strains

- Evolved and ancestral *S. cerevisiae* strains (from Module 1) carrying different pigment expression plasmids

Equipment

- Pipettes: volume needs will vary based on implementation. You will need a P2-20ul, a P20-200ul, and a P200-1000ul or equivalent, as well as a 5ml serological pipette.
- Culture tubes
- Glass beads or plate spreader

Consumables

- YPD + G418 liquid media (at least 30ml experiment)
- YPD + G418 agar plates (at least 4 per experiment)
- Eppendorf tubes
- Sterile swabs, sterile inoculating loops, or sterile inoculating sticks

Chemicals

- FungiCure spray with active ingredient clotrimazole

*Optional*

- *30ºC incubator*
- *Test tube roller drum or shaking platform*
- *Vortex machine*

#### BEFORE THE LAB

1. Plan out how the timing of activities will fit with your class schedule. Yeast grow most robustly at 30ºC. They can be grown at room temperature as well but will grow more slowly. *We've included estimates for the time it'll take for your students' yeast to grow where applicable in italics*.
2. Make FungiCure media. Each competition performed will require 5ml each of a low and a high dose of FungiCure media. In our hands, the following concentrations work well:
    - Low dose: 1:12,800x, 2.25µM (3.88ul of fungicure in 50ml of YPD + G418) inhibits growth the ancestral strains and was the starting concentration for the evolutions.
    - High dose: 1:3200x, 9µM (15.5ul of fungicure in 50ml of YPD + G418) prevents growth of the ancestral strains but not the evolved strains.
    - Very high dose: 1:800x, 36µM (62.1ul in 50ml of YPD + G418) prevents growth of the ancestral strains and several evolved strains, but some evolved strains will grow.

3. Streak evolved and ancestral strains onto YPD + G418 agar media at least 2 days before the intended start of the lab.

**PROTOCOL**

**Day 1:** Inoculate evolved and ancestral strains of yeast into separate tubes of medium.

1. Fill two test tubes with 5ml each of liquid YPD + G418. Label one "evolved" and the other "ancestor".
2. Use a sterile swab, inoculating loop, or inoculating stick to pick a colony of either evolved or ancestral yeast and inoculate it into its respective test tube.
3. Allow the yeast in these tubes to grow until you can no longer see through the liquid media. *When growing at 30$^o$C in a roller drum or shaking platform, this will take 1-2 days. When growing on a bench top at room temperature without shaking or rolling it will take 2-3 days. Yeast can be left longer than these amounts of time (up to a week) without worry.*

**Day 2:** Mix evolved and ancestral strains in media with or without FungiCure.

1. Fill one test tube each with 5ml of the following three media: YPD + G418; YPD + G418 + low dose of FungiCure; YPD + G418 + high dose of FungiCure; YPD + G418 + very high dose of FungiCure. Label these tubes with the media type used.
2. Examine the cultures you inoculated on Day 1. If yeast have settled at the bottom of the tube (pelleted), gently shake the tube until they are completely resuspended. The liquid sample of yeast (culture) should be dense enough that you cannot see through it, and the two cultures should be comparable in density.
3. Remove 20ul of each culture and mix them together in a single eppendorf tube. Make sure to mix well by vortexing or pipetting up and down several times.
4. Add 5ul of mixed culture to each of the three test tubes you inoculated. *Incubate these as on Day 1.*
5. Prepare a 1:10,000 dilution of the remaining mixed yeast culture via serial dilution. Fill two eppendorf tubes with 990ul water and label them "dilution 1" and "dilution 2". In the tube labeled "dilution 1", add 10ul of mixed culture and vortex for 5 seconds or invert 10 times to mix. Transfer 10ul from "dilution 1" to "dilution 2" and vortex for 5 seconds to mix.
6. Use a sterile plate spreader or glass beads to spread 150ul from "dilution 2" onto a YPD + G418 agar plate. *Allow this plate to grow until colonies have formed and their colors can be clearly distinguished (2-3 days at 30$^o$C; 3-4 days at room temperature). Plates can be left at 30$^o$C for 5 days or room temperature for a week without worry.*

**Day 3:** Plate mixed cultures onto YPD + G418 agar media.

1. Examine the cultures you inoculated on Day 2. If yeast have settled at the bottom of the tube (pelleted), gently shake the tube until they are completely resuspended. The liquid sample of yeast (culture) should be dense enough that you cannot see through it, though the culture grown in the highest dose of FungiCure may appear less dense than the others.
2. Prepare a 1:10,000 dilution of each mixed yeast culture via serial dilution. For instance, for the YPD + G418 media condition, fill two eppendorf tubes with 990ul water and label them "YPD + G418 dilution 1" and "YPD + G418 dilution 2". In the tube labeled "YPD + G418 dilution 1", add 10ul of mixed culture and vortex for 5 seconds or invert 10 times to mix. Transfer 10ul from "YPD + G418 dilution 1" to "YPD + G418 dilution 2" and vortex for 5 seconds to mix.
3. Use a sterile plate spreader or glass beads to spread 150ul from "YPD + G418 dilution 2" onto a YPD + G418 agar plate. *Allow this plate to grow until colonies have formed and their colors can be clearly distinguished (2-3 days at 30ºC; 3-4 days at room temperature).*

**Day 4:** Calculate ratio of colors among CFUs as a proxy for relative fitness.

1. Count the number of colonies on each plate onto which you spread yeast culture.
2. Consider the following questions.

#### QUESTIONS

1. Do you see an equivalent number of colonies of each color on the plate you made on Day 2?
2. Does the color ratio differ across the three media conditions you plated from on Day 3?
3. Which strain produced the most colonies after competition in YPD + G418?
4. Which strain produced the most colonies after competition in YPD + G418 + a high dose of FungiCure?

#### EXTENSION QUESTIONS

1. How do your results compare to those of other groups?
2. Do yeast with a high fitness in one environment (presence of FungiCure) always have a high fitness in other environments (absence of FungiCure)?
3. Do all yeast adapted to the same environment (evolved in the same dose of FungiCure) have the same fitness in that environment?
