## Supplemental Text 6 for "yEvo: a modular eukaryotic genetics and evolution research experience for high school students"

### Azole Resistance Module 5
### Metabolic Tradeoffs

#### GOALS

1. Observe metabolic phenotypes by growing yeast on different media.
2. Understand that increased fitness in one environment may come at the expense of decreased fitness in a different environmental condition.
3. Measure the frequency of metabolic tradeoffs in azole-resistant yeast.

#### OVERVIEW

Yeast, like all living organisms, need a source of carbon in their diet. Yeast are most commonly fed a sugar called dextrose, which is the D in YPD growth medium. The evolution experiments that produced the azole-resistant strains you will work with in this module used YPD medium. Yeast can process dextrose through cellular respiration and through fermentation. Some mutations can prevent yeast from undergoing cellular respiration, such as loss of their mitochondrial genome. These mutations prevent growth on non-fermentable carbon sources (e.g. ethanol or glycerol) and lead to slower growth on fermentable carbon sources like dextrose. Because of this slower growth phenotype, respiratory-deficient strains produce smaller colonies and are referred to as "petite mutants". It may seem strange that an evolution experiment would produce yeast that grow more slowly. Shouldn't all mutations that increase fitness lead to faster growth? This paradox is an example of an evolutionary trade-off.

Many of the yeast we've isolated from your experiments are petite due to a loss of mitochondrial DNA. Previous work from other labs has demonstrated that petite mutants tend to have a higher resistance to azole drugs, such as the active ingredient in FungiCure (clotrimazole). In this lab, you will estimate the frequency of petite mutations in your experiments. To do this, you will isolate individual cells from your culture by plating them at low density on YPD, so that each cell can grow up and form a genetically-identical colony. You will then transfer these to a YPG plate (glycerol instead of dextrose) and see which ones are capable of utilizing this carbon source.

#### GLOSSARY

- Cellular respiration: metabolic process to release energy from carbon compounds that occurs in the mitochondria and requires oxygen
- Clotrimazole: An azole antifungal. Inhibits synthesis of ergosterol, a key membrane component and the fungal equivalent of cholesterol. Clotrimazole is the active ingredient in the FungiCure spray used in this experiment.
- Fermentation: metabolic process to release energy from carbon compounds that does not require oxygen or mitochondria

- Fitness: A measure of an individual's reproductive success.
- Petite: small colonies produced by yeast strains that are deficient in cellular respiration
- Selection pressure: An environmental condition that favors some genotypes in a population over others.
- YPD: Yeast Extract, Peptone, and Dextrose; a standard rich yeast medium named for its three ingredients. Also referred to as YEPD.
- YPG: Yeast Extract, Peptone, and Glycerol; a rich yeast medium that contains a non-fermentable carbon source (glycerol). Also referred to as YEPG.

#### MATERIALS AND EQUIPMENT

Yeast strains

- Evolved and ancestral *S. cerevisiae* strains (from Module 1)

Equipment

- Sharpies and rulers

Consumables

- YPD agar plates (1 per strain tested)
- YPG agar plates (1 per strain tested)
- Sterile swabs, sterile inoculating loops, or sterile inoculating sticks

*Optional*

- *30°C incubator*

#### BEFORE THE LAB

1. Plan out how the timing of activities will fit with your class schedule. Yeast grow most robustly at 30°C. They can be grown at room temperature as well but will grow more slowly. *We've included estimates for the time it'll take for your students' yeast to grow where applicable in italics*.
2. Streak evolved and ancestral strains onto YPD + G418 agar media at least 2 days before the intended start of the lab.

**PROTOCOL**

**Day 1:** Grow evolved and ancestral strains of yeast on permissive YPD media.

1. Streak yeast from one of your ancestral and one or more of your evolved populations onto a YPD plate.
2. Let them grow until you can clearly distinguish colonies on the plate. *When growing at 30ºC, this will take 1-2 days. When growing on a bench top at room temperature it will take 2-3 days. Yeast can be left longer than these amounts of time (up to a week) without worry.*

**Day 2:** Grow evolved and ancestral strains of yeast on selective YPG media.

1. Use a sharpie and a ruler to draw a grid on the back of a YPG (glycerol) plate. The grid should have at least 10 evenly-sized boxes. See **Figure 1A** below for an example with 12 evenly-sized boxes.
2. Use a sterile utensil to pick 10 colonies and "patch" them onto your YPG plate by spreading them evenly within the outline of one of the boxes you drew. You don't need to cover the entire box—in fact it's best to leave a little space at the edges so cells from one box don't intrude on a neighboring box. See **Figure 1B** below for an example.

   *To get a good estimate of frequency, it's important to come up with a scheme that reduces experimental bias. This could be "I chose the last 10 colonies from my streak", or "half my colonies were large and half were small, so I picked 5 large and 5 small colonies".*
3. Record the phenotype of each patched colony (was it larger or smaller than other colonies on the plate?).
4. Allow yeast to grow on YPG plate for 2-7 days.

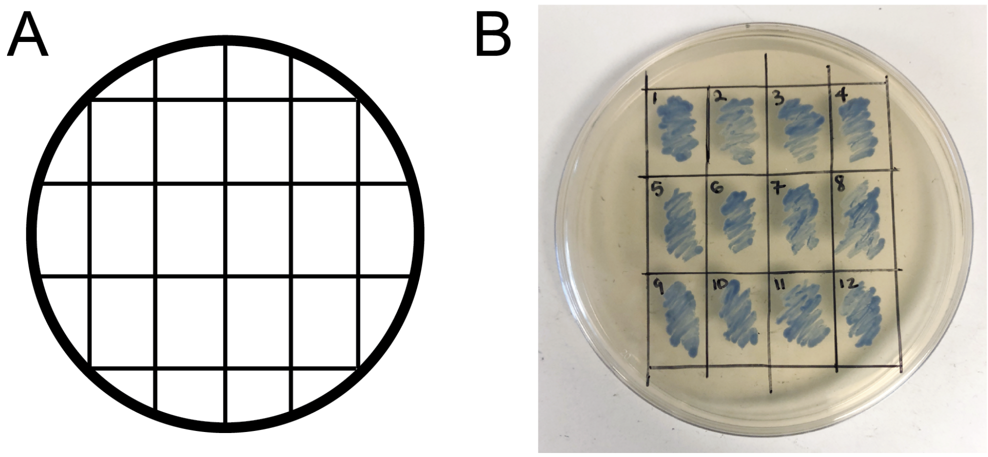


**Figure 1: Patching colonies.** (A) Example of a 12-box grid on a plate. (B) Colonies of a blue yeast strain patched into 12 boxes on a YPD plate. Note that space is left around each patch.

**Day 3:** Record phenotypes from growth on YPG.

1. After you can clearly see colonies, answer the following questions.

#### QUESTIONS

1. What fraction of patched colonies grew on YPG?
2. What fraction of patched colonies were petite?
3. Do phenotypes on YPD plates correlate with any phenotypes you observed on YPG plates?

#### EXTENSION QUESTIONS

1. How would your results differ if you had used a YPG instead of a YPD plate on Day 1?
2. Do you see variability in petite frequency between your evolved replicates? How about between your replicates and your classmates?
3. Does the type of media in which you evolved your yeast impact the types of azole resistance mutations that arose during your experiment?
4. Why would a mutation that makes yeast grow slowly be helpful in some conditions?
5. Why would losing cellular respiration be helpful in dealing with FungiCure?
